# Oral prodrug of a novel glutathione surrogate reverses metabolic dysregulation and attenuates neurodegenerative process in APP/PS1 mice

**DOI:** 10.1101/2025.01.15.633247

**Authors:** Swetha Pavani Rao, Aminat O. Imam-Fulani, Wei Xie, Samuel Phillip, Krishna Chennavajula, Erin B. Lind, Ying Zhang, Robert Vince, Michael K. Lee, Swati S. More

## Abstract

Glycation-induced oxidative stress underlies the numerous metabolic ravages of Alzheimer’s disease (AD). Reduced glutathione levels in AD lead to increased oxidative stress, including glycation-induced pathology. Previously, we showed that the accumulation of reactive 1,2-dicarbonyls such as methylglyoxal, the major precursor of non-enzymatic glycation products, was reduced by the increased function of GSH-dependent glyoxalase-1 enzyme in the brain. In this two-pronged study, we evaluate the therapeutic efficacy of an orally bioavailable prodrug of our lead glyoxalase substrate, pro-ψ-GSH, for the first time in a transgenic Alzheimer’s disease mouse model. This prodrug delivers pharmacodynamically relevant brain concentrations of ψ-GSH upon oral delivery. Chronic oral dosing of pro-ψ-GSH effectively reverses the cognitive decline observed in the APP/PS1 mouse model. The prodrug successfully mirrors the robust effects of the parent drug i.e., reducing amyloid pathology, glycation stress, neuroinflammation, and the resultant neurodegeneration in these mice. We also report the first metabolomics study of such a treatment, which yields key biomarkers linked to the reversal of AD-related metabolic dysregulation. Collectively, this study establishes pro-ψ-GSH as a viable, disease-modifying therapy for AD and paves the way for further preclinical advancement of such therapeutics. Metabolomic signatures identified could prove beneficial in the development of treatment-specific clinically translatable biomarkers.

**ABSTRACT GRAPHIC:** 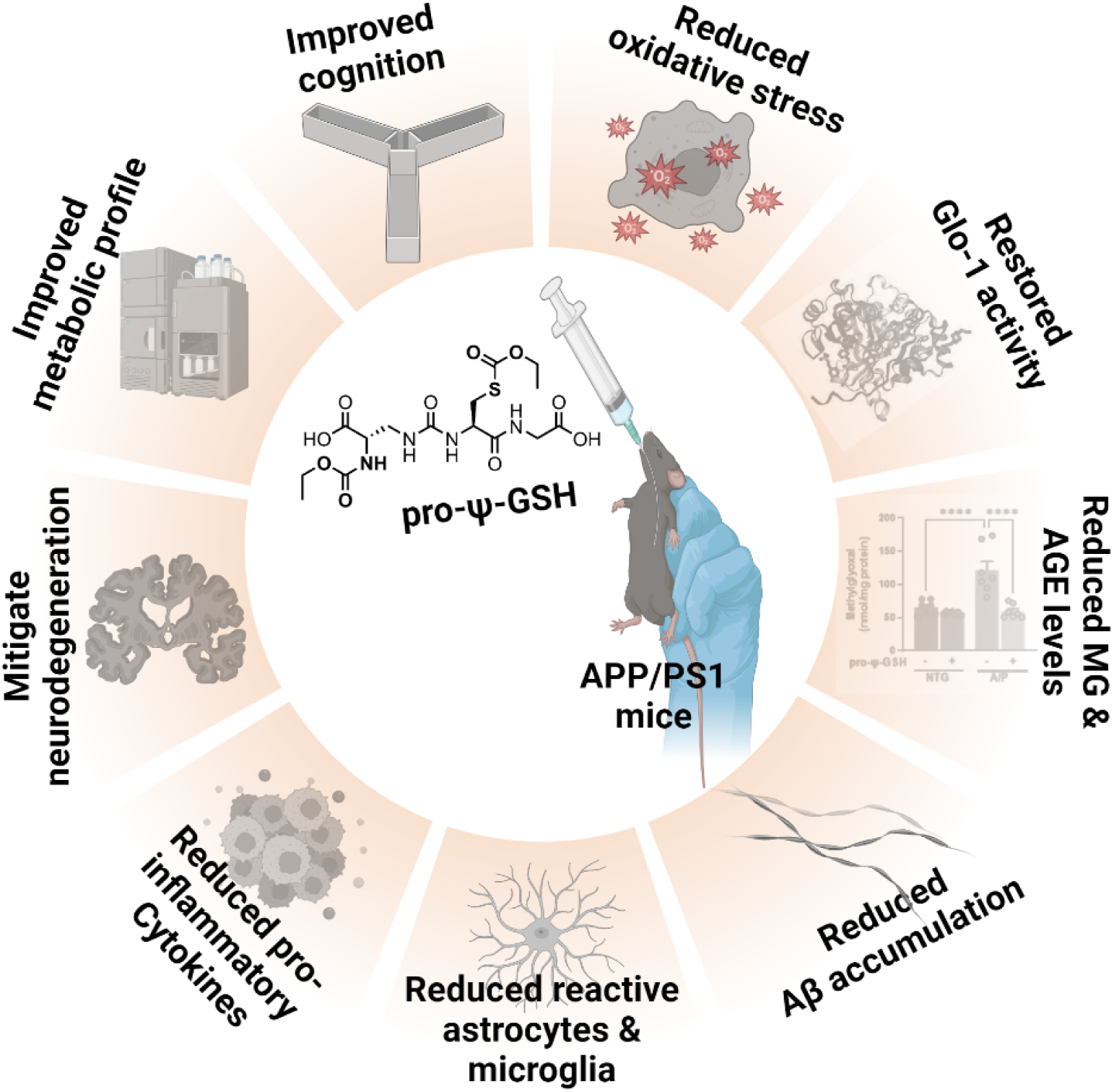

---

Alzheimer’s disease (AD) and related dementias are chronic debilitating conditions that contribute significantly toward cognitive impairment in elderly. AD is a progressive neurodegenerative disease that accounts for 60-70% of neurodegenerative dementia cases.^1^ Current pharmacotherapy has been limited to addressing the neurological symptoms, without addressing the underlying disease biology. Recently approved antibody-based therapies targeting amyloid aggregates show modest slowing of disease progression, offering marginal cognitive benefits, however, with significant side effects.^2, 3^ Since AD is a multifactorial disorder,^4^ the development of additional disease-modifying therapies aimed at multiple target proteins contributing to its etiology is warranted.

The key neuropathological hallmarks of AD are extracellular deposits of β-amyloid peptide (Aβ), intracellular aggregates of phosphorylated forms of tau, and progressive neurodegeneration.^5^ Studies show that AD pathology is invariably associated with increased oxidative stress and brain inflammation. One of the consistent signs of oxidative stress in AD is the decreased levels of reduced glutathione (GSH), indicating increased oxidative stress and inflammation in AD.^6, 7^ Specifically, in addition to being the major antioxidant in the brain, GSH is required as a co-factor for many antioxidant enzymes, including glyoxalase-1 (Glo-1). Glo-1 metabolizes reactive dicarbonyls (e.g. methylglyoxal, MG) resulting from normal glycolysis and oxidization of sugars (Figure 1A).^8^ If not neutralized by Glo-1, MG forms irreversible adducts with basic amino acids, arginine, and lysine present in proteins, resulting in the formation of covalent adducts in the form of hydroimidazolone (MG-H1), argpyrimidine, Nε-carboxyethyl-lysine (CEL), etc., referred to as advanced glycation end products (AGEs).^9, 10^ Such AGEs are highly pro-inflammatory and oxidant substances that are implicated in diabetes, atherosclerosis, and other metabolic disorders.^11, 12^ In AD, increased AGE levels and brain inflammation are associated with reduced Glo-1 activity.^13^ Similarly, we showed that in the APP/PS1 model, amyloid pathology is sufficient to cause a decrease in Glo-1 activity and an increase in MG levels.^14^

**Figure 1.**
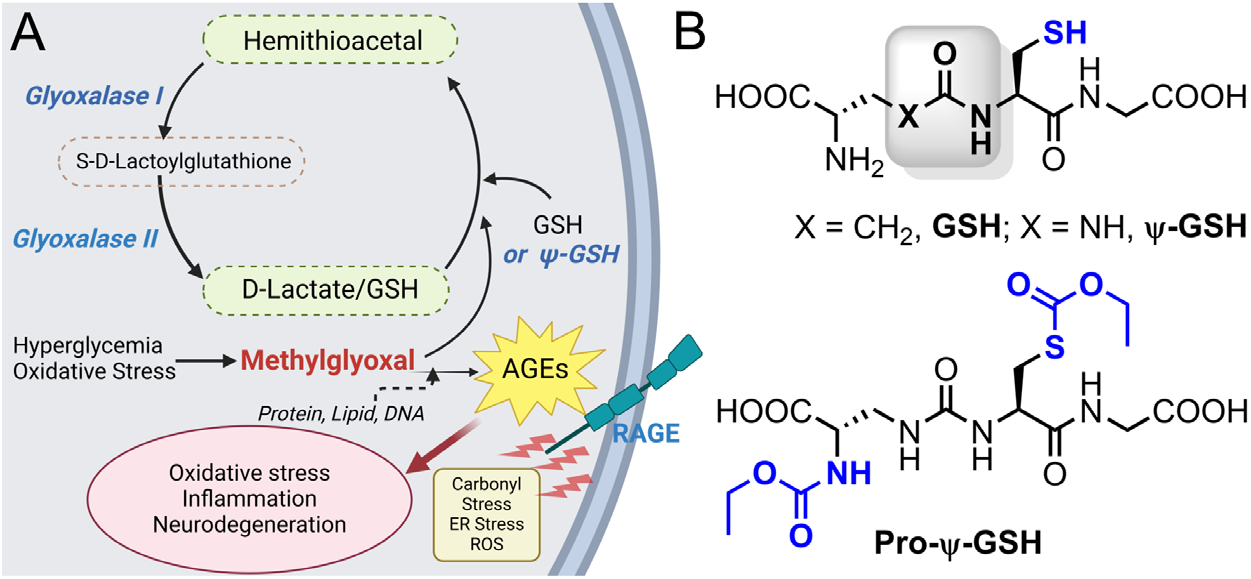
(A) Glyoxalase enzymatic pathway involved in detoxification of methylglyoxal. Formation of advanced glycation end products (AGE) by methylglyoxal with biological macromolecules results in oxidative stress and inflammatory cascade. (B) Chemical structures of the glyoxalase substrate glutathione (GSH), ψ-GSH, and the prodrug, pro-ψ-GSH.

While it is attractive to use GSH supplementation for AD,^7^ the instability of GSH to γ-glutamyl transpeptidase (GGT) renders sufficient brain supplementation impossible. To address this, we have developed a ureide peptidomimetic of GSH, ψ-GSH, which resists GGT-mediated cleavage.^15^ Given intraperitoneally, ψ-GSH counters oxidative stress, cognitive decline, and neuropathology in the APP/PS1 mouse model — even when given after the onset of symptoms.^14, 16^ Because oral ψ-GSH was ineffective in achieving these end points, we developed prodrugs of ψ-GSH that could effectively deliver the parent compound to the brain.^17^ The site of metabolic instability and poor bioavailability of ψ-GSH was the free sulfhydryl group, which renders it susceptible to oxidative metabolism. Through our preliminary pharmacokinetic study, S-ethylcarbamoyl ψ-GSH (pro-ψ-GSH, Figure 1B) emerged as the lead prodrug that could effectively convert into parent compound ψ-GSH, offered better stability in the gastrointestinal tract, and improved plasma stability to deliver ψ-GSH across the blood-brain barrier.^17^ Oral administration of pro-ψ-GSH to acute, AD-like mouse model generated by intracerebroventricular (i.c.v) administration of amyloid-β peptides demonstrated neuroprotective activity and effective reversal of working memory deficits. The prodrug countered oxidative and inflammatory consequences of amyloid aggregates by restoring GSH levels, akin to intraperitoneal administration of the parent ψ-GSH.^17^ The results of this study provided a rational basis for conducting a thorough preclinical evaluation of pro-ψ-GSH in a transgenic AD animal model that better represents human AD pathology.

Herein, we performed a preclinical evaluation of pro-ψ-GSH in the APP/PS1 transgenic mouse model of AD. We show that the prodrug was able to attenuate cognitive deficits and AD-like pathology in these mice. Biochemical and immunohistochemical analysis demonstrated the effect of the treatment on AD pathology, glycation-induced oxidative and inflammatory changes, and neurodegeneration. Metabolic analysis of the hippocampal region identified key signatures that could be useful in future translational studies. These findings thus establish mitigation of glycation-induced oxidative stress to be a lucrative multipronged therapeutic strategy that is needed to achieve true disease modification in AD. Consequently, a warrant is thus established for future preclinical advancement of pro-ψ-GSH therapy in animal models displaying both amyloid and tau pathologies.

## RESULTS

### Oral pro-ψ-GSH treatment improves cognitive function in APP/PS1 mice

Based on the previous promising results in acute Aβ toxicity model,^17^ we conducted a chronic efficacy evaluation of pro-ψ-GSH in the APP/PS1 transgenic mouse model of AD that better recapitulates many aspects of human AD pathology. Oral treatment of pro-ψ-GSH (250 mg/kg) was initiated in 8-9-month old APP/PS1 mice and continued for 12 weeks before assessment of mice for cognitive function, biochemical evaluation of brain tissue, and neuropathology. We performed a battery of cognitive behavioral tests, such as the Y-maze, Barnes maze, and novel object recognition test, to understand the impact of the treatment on crucial cognitive functions, including spatial working memory, spatial learning, and reference memory (Figure 2).^18^

**Figure 2.**
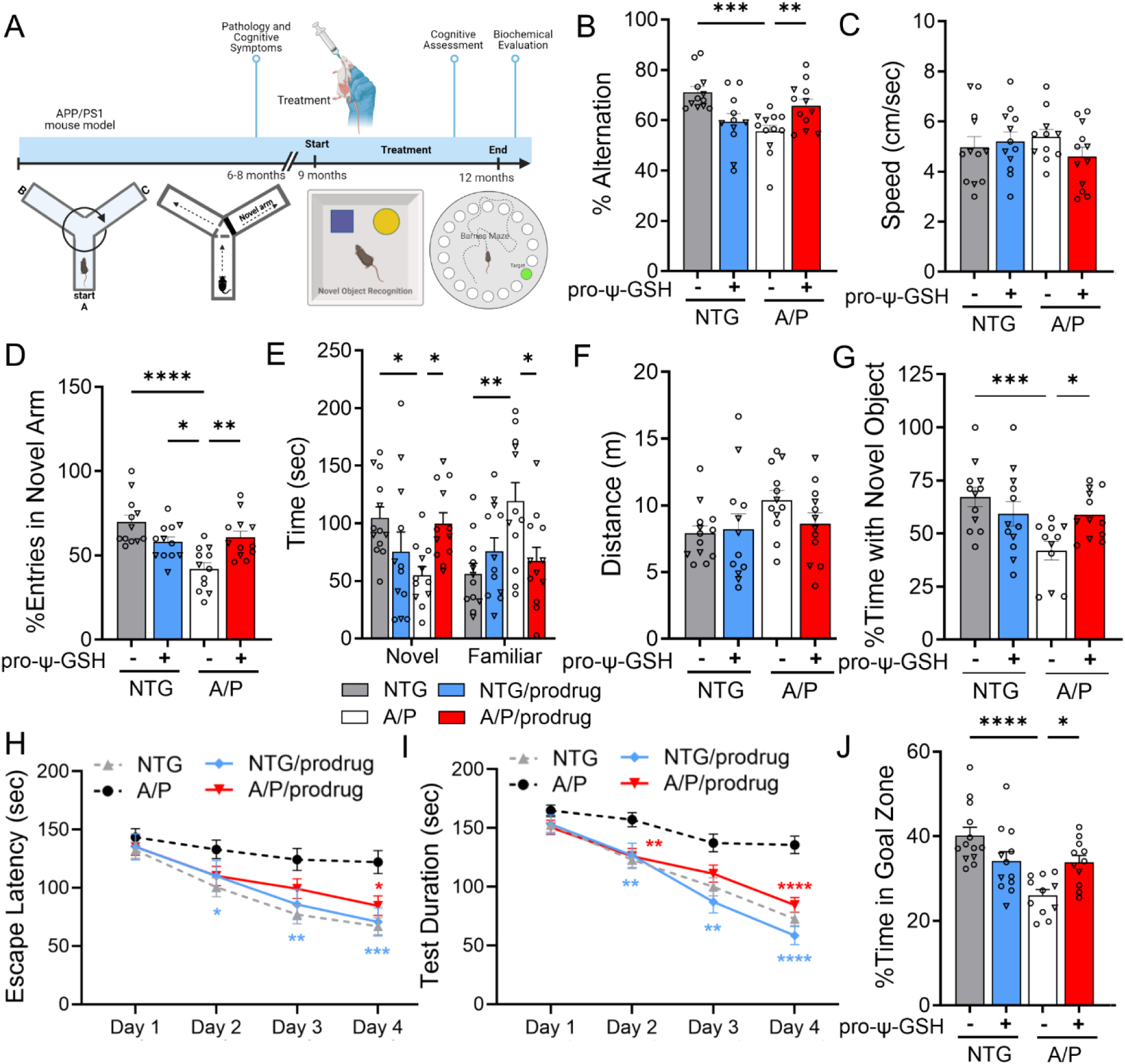
Oral administration of pro-ψ-GSH in symptomatic APP/PS1 mice reverses cognitive impairment. (A) Schematic of the experimental timeline in aged APP/PS1 (A/P) mice. (B) Percent spontaneous alternations in Y-maze of animals treated for 12 weeks. Significant improvement in the alternation behavior was observed between prodrug and saline treated APP/PS1 mice. (C) Average speed of the mice during the Y-maze alternation test were unaffected by the genotype and the prodrug treatment. (D-F) In Y-maze spatial recognition test, the prodrug treated APP/PS1 group spent more time in the previously unexplored (novel) arm compared to the familiar arm, as evident from significantly higher number of entries (D) in the novel arm, and corollary, more time spent (E) in the novel arm. The total distance traveled (F) was unchanged within the genotypes and treatment groups. (G) In novel object recognition test, prodrug-treated APP/PS1 mice exhibited higher exploratory behavior with the novel object compared to corresponding vehicle controls. (H-I) Escape latency and test duration for identification of target hole during the training period in Barnes maze. Prodrug-treated APP/PS1 mice learned to find the escape hole faster than saline-treated APP/PS1 mice over the test duration. (J) Proportion of time spent in the target quadrant over rest of the maze area during the probe trial conducted 24 h after the last training session. Oral pro-ψ–GSH group performed significantly better in the probe trial, and spent larger proportion of total time in the target zone compared to the controls. Circles are males and triangles are females. Data are represented as mean ± SEM. One-way ANOVA or two-way ANOVA (for E) followed by Tukey’s or Holm-Sidak’s post-hoc multiple comparisons test *p<0.05, **p<0.01, ***p<0.001, and ****p<0.0001.

### Y-maze Spontaneous Alternation test

The Y-maze spontaneous test is widely used for evaluating short-term memory in mice.^19^ This test depends on the innate curiosity of mice to explore a novel arm, resulting in prominent alternation behavior, which is compromised due to the deposition of Aβ in hippocampus, prefrontal cortex regions. In this study, such spatial working memory deficits were apparent in vehicle-treated symptomatic APP/PS1 cohort; specifically, approximately 22% reduction in alternation rate was observed in saline-treated APP/PS1 mice compared to corresponding age-matched wild-type (NTG) mice (NTG-veh: A/P-veh; 71.17 ± 2.23%, 55.54 ± 2.50, respectively, Figure 2B). The working memory deficits were reversed after pro-ψ-GSH treatment (A/P-pro-ψ-GSH; 65.78 ± 2.64%). The total distance traveled and time taken to complete the test (Figure 2C) were unchanged among the treatment groups. This confirmed the lack of motor dysfunction or physical incapacitation of mice influencing the observed trend. Additionally, pro-ψ-GSH treated wild type mice showed behavior similar to the saline controls.

### Y-Maze Spatial recognition test

This Y-maze test is adapted to evaluate changes related to spatial recognition memory.^20^ Spatial recognition memory was assessed based on the natural tendency of mice to preferentially explore novel areas over familiar environments in a two trial Y-maze test. The number of entries in the novel arm and time spent between novel *vs* familiar arms was used to define spatial memory changes. In comparison, among the vehicle-treated NTG and APP/PS1 groups, APP/PS1 mice displayed deficits in spatial recognition (fewer entries in the novel arm) due to impaired hippocampal function. Administration of pro-ψ-GSH resulted in a significant 1.45-fold increase in their interest in exploring the novel arm as calculated from percent entries in the novel arm (NTG-veh: A/P-veh: A/P-pro-ψ-GSH; 69.86 ± 4.17%, 42.06 ± 3.61, 60.79 ± 3.69, respectively; Figure 2D). This was reciprocated in the total time spent in the novel arm (Figure 2E). While the APP/PS1 saline group spent significantly less time in the novel arm, the group spent approximately 77% more time in the familiar arm compared to the NTG-saline controls. Reversal of this behavior was observed by oral pro-ψ-GSH treatment (NTG-veh: A/P-veh: A/P-pro-ψ-GSH; 56.15 ± 9.06%, 119.1 ± 16.2, 67.36 ± 11.7, respectively), signifying the beneficial effects of the treatment. There were no significant differences in the overall activity of mice, which is evident from the unchanged total distance travelled (Figure 2F). The compound did not show any significant observations in the NTG mice.

### Novel Object Recognition Test (NORT)

This test leverages the natural behavior of mice to interact with a novelty over familiarity is the Novel Object Recognition test.^21^ We performed NORT on prodrug treated NTG and APP/PS1 mice in comparison with corresponding saline-treated groups. This test is considered low-stress and efficient, and does not concern the spatial reference memory of the animals. In the training phase, mice are allowed to explore two identical objects, while in the testing phase, one of the two identical objects is replaced by a novel one. Saline and pro-ψ-GSH treated NTG mice showed a clear preference for the novel object exploration as evident from the percent time spent with the novel object (Figure 2G). APP/PS1 mice showed reduced interaction with the novel object, possibly due to lack of recall of the familiar object presented during the initial trial. The preference for the novel object was improved upon pro-ψ-GSH treatment (NTG-veh: A/P-veh: A/P-pro-ψ-GSH; 67.29 ± 4.54%, 41.92 ± 4.33, 59.00 ± 3.23, respectively; Figure 2G), suggesting reversal of recall deficit caused by AD pathology.

### Barnes maze

The use of multiple tests is important to capture the complexity of cognitive processes and ensure that the findings are robust and not an artifact of a single behavioral paradigm.^22^ Hence, we evaluated the spatial memory in APP/PS1 mice using the Barnes maze test, as described in the Methods section, which evaluates long term spatial memory and learning behavior. The escape latency of mice irrespective of genotype and treatment, on the first day of training showed no significant differences. As the training progressed, the NTG-saline treated mice performed better with a significant reduction in the escape latency (Figure 2H) and test duration (Figure 2I), whereas no such improvement was noted in saline-treated APP/PS1 mice. Pro-ψ-GSH treatment of APP/PS1 mice resulted in reduction in the time required to locate the escape hole, showing a learning behavior similar to NTG controls. Specifically, the treatment group on days 3 and 4 showed statistically significant attenuation of escape latency compared to corresponding untreated APP/PS1 mice (Day 3 — NTG-veh: A/P-veh: A/P-pro-ψ-GSH; 77.09 ± 7.98, 124.3 ± 9.49, 99.33 ± 8.39 sec, respectively; Day 4 — NTG-veh: A /P-veh: A/P-pro-ψ-GSH; 66.96 ± 7.26, 122.1 ± 9.78, 84.61 ± 8.42 sec, respectively; Figure 2H). A probe trial conducted 24 hours after the last training trial showed improvement in the retention of learned tasks in the oral pro-ψ-GSH treated group. Time spent in the target quadrant by pro-ψ-GSH treated mice was significantly higher than the untreated transgenic mice, showing improvement in spatial reference memory (NTG-veh: A/P-veh: A/P-pro-ψ-GSH; 40.11 ± 1.90, 26.01 ± 1.37, 33.9 ± 1.60 sec, respectively, Figure 2J). Additionally, the distance traveled by APP/PS1 mice showed a significant reduction of nearly 20%, whereas pro-ψ-GSH treatment significantly mitigated the impaired effects in APP/PS1 mice.

### Oral pro-ψ-GSH restored glyoxalase-1 activity leading to reduction in methylglyoxal levels

Herein, we aimed to investigate whether orally administered pro-ψ-GSH could deliver ψ-GSH at pharmacologically relevant concentrations and restore compromised Glo-1 activity in APP/PS1 mice. To determine this, we first quantitated the levels of ψ-GSH in the brain tissue at the end of the treatment period. Confirming our previous pharmacokinetic evaluation ^14, 17^, pro-ψ-GSH efficiently delivered the parent compound at levels found to be efficacious (Supplementary Figure S1). We then examined the expression of Glo-1 and Glo-2 proteins in mouse brain cortical tissue using western blot analysis and observed no changes in the expression levels in all groups (Figure 3A, B). This confirmed observation from our and other previous studies that Glo-1 expression is maintained even at this advanced stage in the disease.^13, 14, 23^ Consistent with previous results,^14^ we found a significant decrease in endogenous glyoxalase enzyme activity in APP/PS1 mice compared to that in saline-treated NTG mice (Figure 3C) (NTG-veh: A/P-veh; 0.117 ± 0.002, 0.100 ± 0.002, OD_240_/mg protein, respectively). Pro-ψ-GSH treatment was able to supplement the compromised Glo-1 function in in APP/PS1 mice to NTG control levels (A/P-pro-ψ-GSH: 0.113 ± 0.002, ΔOD/min/mg protein). The downstream effects of Glo-1 function supplementation were examined by measurement of the levels of the substrate MG, and the covalent adducts of MG with proteins in the form of AGEs. We found about 1.5-fold increase in the MG levels in APP/PS1 mouse brain compared to the NTG brain tissue (Figure 3D), correlating with the reduced enzymatic function. Pro-ψ-GSH treatment completely normalized MG levels in APP/PS1 mice to those found in NTG mice (NTG-veh: A/P-veh: A/P-pro-ψ-GSH; 65.22 ± 4.10, 121.4 ± 13.6, 59.21 ± 3.91 nmol/mg protein, respectively). Similarly, increased accumulation of AGEs in APP/PS1 brains was significantly reduced by the prodrug (Figure 3E), attesting to delivery of parent ψ-GSH and in vivo engagement of Glo-1 pathway (NTG-veh: A/P-veh: A/P-pro-ψ-GSH; 4.796 ± 0.15, 6.283 ± 0.20, 5.301 ± 0.16 ng/mg protein, respectively). Analysis of NTG mice shows that pro-ψ-GSH treatment did not alter any of these parameters.

**Figure 3.**
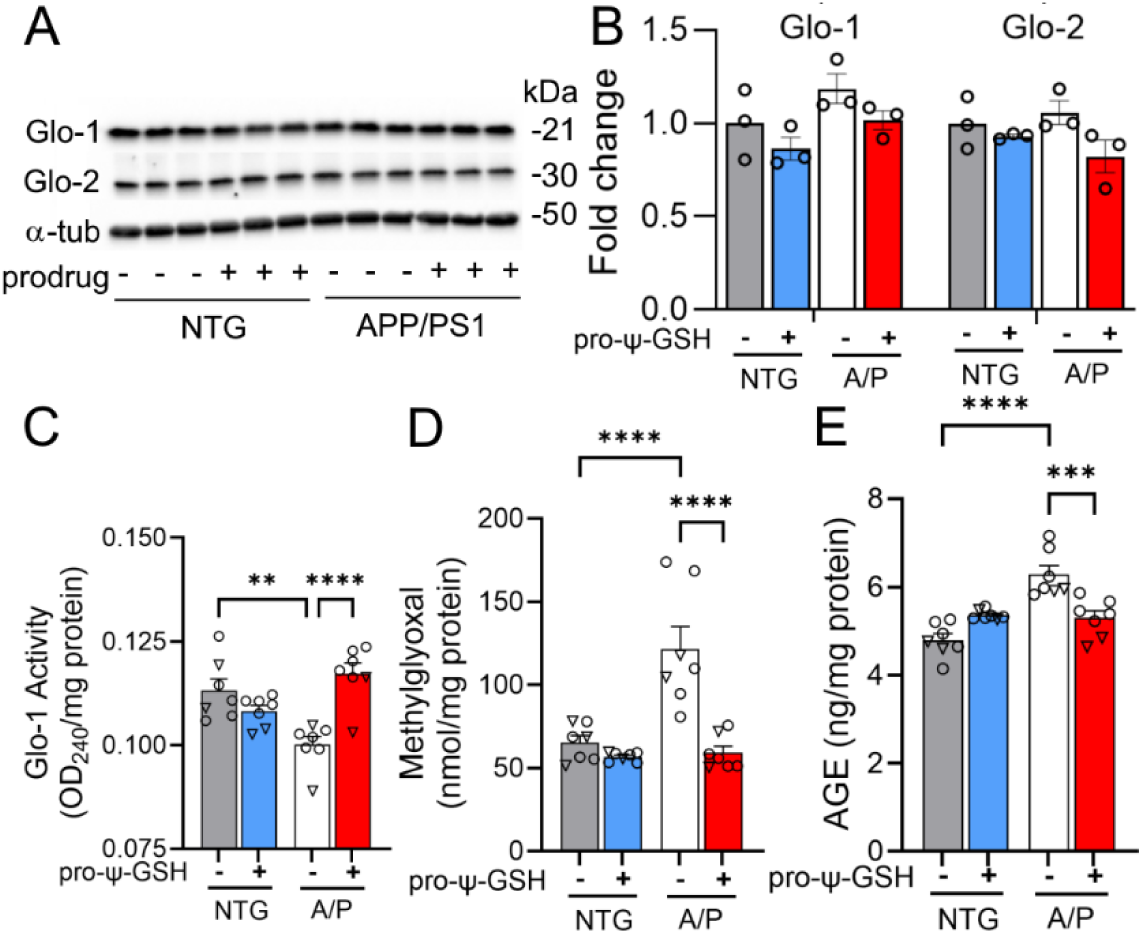
Effect of pro-ψ-GSH supplementation on Glo-1 related pathway markers in aged APP/PS1 mice. (A) Glo-1 and Glo-2 protein expression analyses by western blot and (B) their respective quantification. (C) Glo-1 enzyme activity was significantly compromised in APP/PS1 (A/P) mice compared to the corresponding non-transgenic controls (NTG), which was restored by the oral prodrug treatment. (D) Reduced Glo-1 function resulted in increased MG levels in the APP/PS1-saline group. The MG levels were normalized to NTG-saline group levels by the prodrug. (E) Increased AGEs content, corresponding to higher MG levels, in untreated APP/PS1 mice was reduced significantly by pro-ψ-GSH treatment. Circles are males and triangles are females. Data are presented as the mean ± S.E.M and is representative of two (for A & B) or three (C–E) independent experiments. Pro-ψ-GSH and saline-treated APP/PS1 and NTG groups were compared with one-way ANOVA with Tukey’s post-hoc multiple comparison test for statistical analysis. ** p < 0.01, *** p < 0.001, **** p < 0.0001.

### Reduced oxidative stress in transgenic Alzheimer’s mice treated with pro-ψ-GSH

We tested if oral pro-ψ-GSH, similar to *i.p.* ψ-GSH, can also reverse signs of oxidative stress by examining various indices of oxidative stress such as cellular redox potential (GSH levels, GSH:GSSG ratio), and lipid peroxidation (TBARS assay). We observed that the GSH/GSSG ratio was significantly lower in APP/PS1 mice compared to that in the wild-type controls, confirming increased oxidative stress and reduced antioxidant capacity with AD pathology in the APP/PS1 model (Figures 4A, B). However, prodrug treatment corrected the GSH/GSSG ratio, thereby reducing oxidative stress by restoring GSH deficits (NTG-veh: A/P-veh: A/P-pro-ψ-GSH; 0.487 ± 0.06, 0.296 ± 0.03, 0.491 ± 0.04, respectively). Indeed, the levels of reduced GSH were restored to NTG levels in the pro-ψ-GSH treated mice. These results were further substantiated by quantification of lipid peroxidation using the TBARS assay (Figure 4C). Consistent with the results of the GSH assay, we observed elevated malondialdehyde levels in untreated transgenic mice, which were normalized to control levels by pro-ψ-GSH treatment in APP/PS1 mice (NTG-veh: A/P-veh: A/P-pro-ψ-GSH; 5.301 ± 0.59, 8.764 ± 0.49, 5.841 ± 0.85 μM/mg protein, respectively).

**Figure 4.**
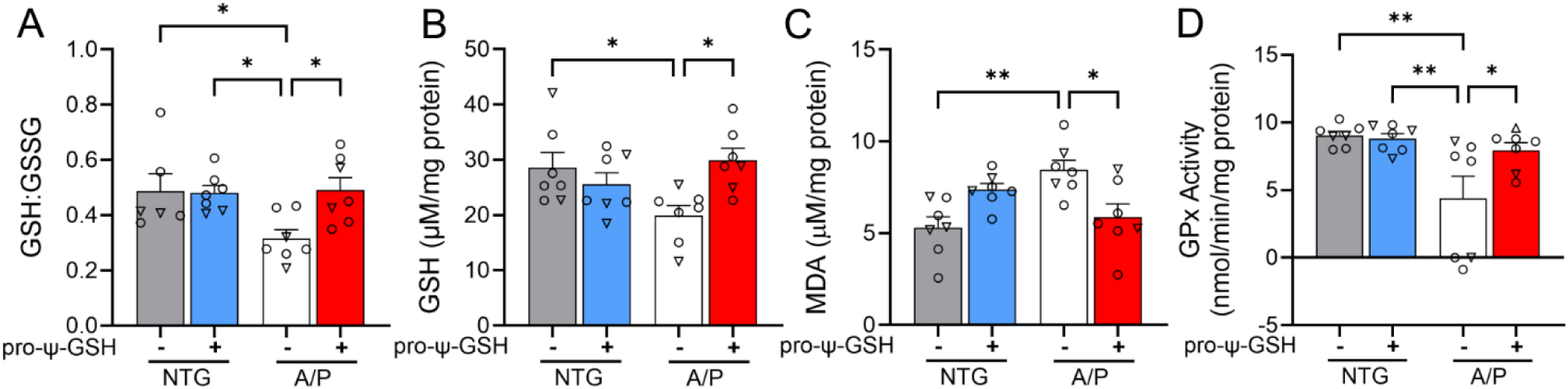
Mitigation of increased oxidative stress by pro-ψ-GSH treatment in APP/PS1 mice brain. (A-B) Improved redox ratio (GSH/GSSG, A) was apparent in pro-ψ-GSH treated APP/PS1 mice. Quantification of reduced GSH (B) showed lower cytoplasmic GSH content in the APP/PS1 brains compared to NTG mice, which were restored by the prodrug treatment. (C) Lipid peroxidation levels as calculated from the levels of MDA were attenuated by the prodrug in the APP/PS1 group. (D) Compromised GPx activity in aged APP/PS1 saline group was restored to NTG levels by pro-ψ-GSH. Circles are males and triangles are females. Data are shown as the mean ± SEM. Data are representative of three independent experiments. Statistical significance was examined by a one-way ANOVA with Tukey’s or Sidak’s post-hoc multiple comparison test. * p < 0.05, ** p < 0.01.

Glutathione peroxidase (GPx) is a selenium containing antioxidant enzyme present in both the cytosol and the mitochondrial matrix. Its role includes facilitating detoxification of hydrogen peroxide to water, and lipid peroxides to corresponding lipid alcohols.^24^ Increased lipid peroxidation in the AD brain has been linked to crippled GPx enzymatic pathway.^25^ Due to its dependence on GSH levels, it is expected that delivery of ψ-GSH via its prodrug form could substitute for GSH in this enzymatic reaction. We observed that GPx activity in APP/PS1 mice was approximately half of that in NTG controls (Figure 4D). Reduced GPx activity was restored by pro-ψ-GSH treatment of APP/PS1 mice to levels found in NTG controls (NTG-veh: A/P-veh: A/P-pro-ψ-GSH; 9.037 ± 0.30, 4.369 ± 1.66, 7.942 ± 0.56 nmol/min/mg protein, respectively). The NTG-pro-ψ-GSH group did not exhibit any changes in activity.

### Oral pro-ψ-GSH treatment reduced brain amyloid pathology in APP/PS1 mice

The APP/PS1 transgenic model used here develops amyloid pathology, a pathological hallmark of AD,^1^ in the forebrain areas at 6 to 7 months of age.^16^ Thus, we asked whether the oral administration of pro-ψ-GSH after the onset of amyloid pathology at ∼9 months of age can attenuate further progression of amyloid pathology at 12 months of age in the APP/PS1 model.

Immunohistochemical analysis of Aβ plaques by 4G8 antibody in saline-treated APP/PS1 mice displayed severe Aβ pathology in the cortex and hippocampal regions (Figures 5A-C). The APP/PS1 group treated with oral pro-ψ-GSH for the prior 3 months exhibited a substantial reduction in the Aβ plaque burden compared to the untreated APP/PS1 group. Quantitative analysis of the S1BF region of the cortex and the hippocampus showed that the brain area covered by 4G8 immunoreactivity was significantly lower in the pro-ψ-GSH treated group compared to the corresponding saline-treated APP/PS1 mice [Cortex (Figure 5B) — A/P-veh: A/P-pro-ψ-GSH; 38.15 ± 4.27%, 22.76 ± 2.69%, respectively, p < 0.01; Hippocampus (Figure 5C) — A/P-veh: A/P-pro-ψ-GSH; 29.98 ± 4.45%, 16.66 ± 1.66%, respectively, p < 0.05]. We further confirmed the neuropathological findings through independent analysis of the brain tissues by an Aβ ELISA assay to determine the effect of the treatment on cortical levels of soluble and insoluble Aβ_1-42_ (Figure 5D-E). The levels of insoluble Aβ_1-42_ were markedly reduced in pro-ψ-GSH treated APP/PS1 mice compared to saline-control APP/PS1 littermates (NTG-veh: A/P-veh: A/P-pro-ψ-GSH; 11.43 ± 0.66, 2887 ± 213, 1707 ± 116 ng/mg protein, respectively). The effect of the treatment on soluble Aβ_1-42_, however, did not reach statistical significance (NTG-veh: A/P-veh: A/P-pro-ψ-GSH; 56.26 ± 3.19, 814.6 ± 75.6, 949.1 ± 66.0 ng/mg protein, respectively).

**Figure 5.**
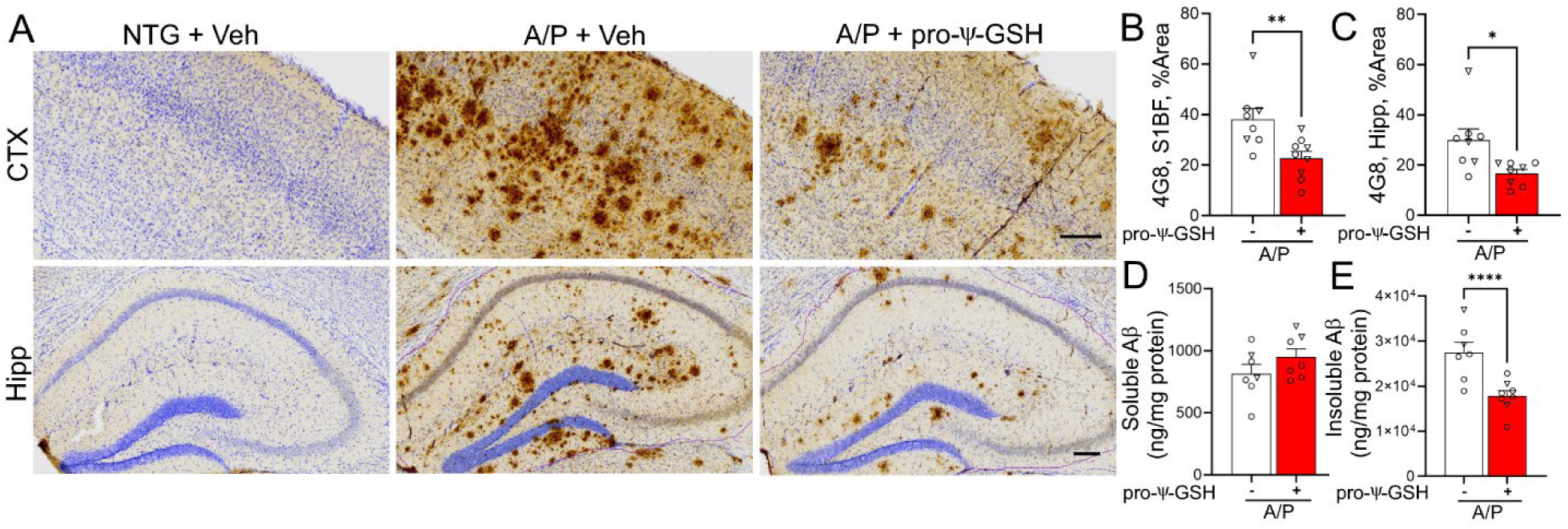
Oral pro-ψ-GSH effectively reduces Aβ burden in symptomatic APP/PS1 mice. (A) Representative images of Aβ immunoreactivity (using 4G8 antibody) in the S1BF cortex (CTX) and hippocampus (Hipp) from non-Tg littermates (NTG) and APPswe/PS1ΔE9 Tg mice (A/P). Animals were treated with either Vehicle (Veh) or pro-ψ-GSH for 3 months prior to tissue collection. Scale bar, 100 µm. (B, C) Quantitative analysis of amyloid pathology was done by determining the percent area covered by 4G8 immunoreactivity in S1BF (B) and hippocampus (C). Compared to the vehicle (Veh) treated mice, pro-ψ-GSH-treated mice showed significantly lower 4G8 immunoreactivity (n = 8 APP/PS1 + Saline; n = 9 APP/PS1 + pro-ψ-GSH). There was no Aβ immunoreactivity in NTG mice. (D, E) ELISA assay quantitation of the β-amyloid load in A/P mice. Levels of insoluble Aβx–42 levels (E) in the brain homogenate of A/P mice treated with pro-ψ-GSH were reduced significantly compared to the Veh treated A/P mice. Circles are males and triangles are females. Unpaired Student’s t-test was used for all statistical comparisons of saline and ψ-GSH-treated A/P cohorts. For comparisons between saline and pro-ψ-GSH-treated APP/PS1 groups, a one-way ANOVA with Tukey’s or Sidak’s post-hoc multiple comparison test was performed for statistical analysis. * p < 0.05, ** p < 0.01, *** p < 0.005. * p < 0.05, ** p < 0.01, *** p < 0.005.

### Oral pro-ψ-GSH mitigates neuroinflammatory stress in symptomatic APP/PS1 mice

Amyloid pathology is associated with higher astrogliosis and microglial activation, leading to overexpression of pro-inflammatory cytokines, and secretory inflammatory factors that ultimately damage neurons.^14^ Because oral pro-ψ-GSH reduces amyloid pathology and MG/AGE levels, we examined if oral pro-ψ-GSH also attenuates inflammatory activation of glial cells in the APP/PS1 model.

As expected, the presence of Aβ pathology in saline-treated APP/PS1 is associated with significant increase in GFAP staining, indicating activation of astrocytes in the cortex and hippocampus compared to the non-transgenic control mice (Figure 6A, B). Treatment with pro-ψ-GSH resulted in significant attenuation of GFAP immunoreactivity (NTG-veh: A/P-veh: A/P-pro-ψ-GSH; 7.960 ± 1.67%, 24.24 ± 2.67%, 16.07 ± 1.80%, respectively). Although a trend toward astrocyte activation was apparent in the hippocampal region of the APP/PS1-vehicle group compared to NTG-vehicle, it did not reach statistical significance (Supplementary Figure S2). Analysis of cortical sections immunostained for microglia using Iba-1 antibody also shows a significant increase in Iba-1 immunoreactivity in saline-treated APP/PS1 cohort (Figure 6C, D). As with astrogliosis, pro-ψ-GSH treatment significantly reduces Iba-1 immunoreactivity in APP/PS1 mice (NTG-veh: A/P-veh: A/P-pro-ψ-GSH; 3.271 ± 0.33%, 9.099 ± 0.74%, 5.082 ± 0.35%, respectively). Treatment of non-transgenic mice with pro-ψ-GSH did not have any significant effect on the level of GFAP and Iba-1 immunoreactivity in both cortex and hippocampus (Figure 6 and Supplementary Figure S3).

**Figure 6.**
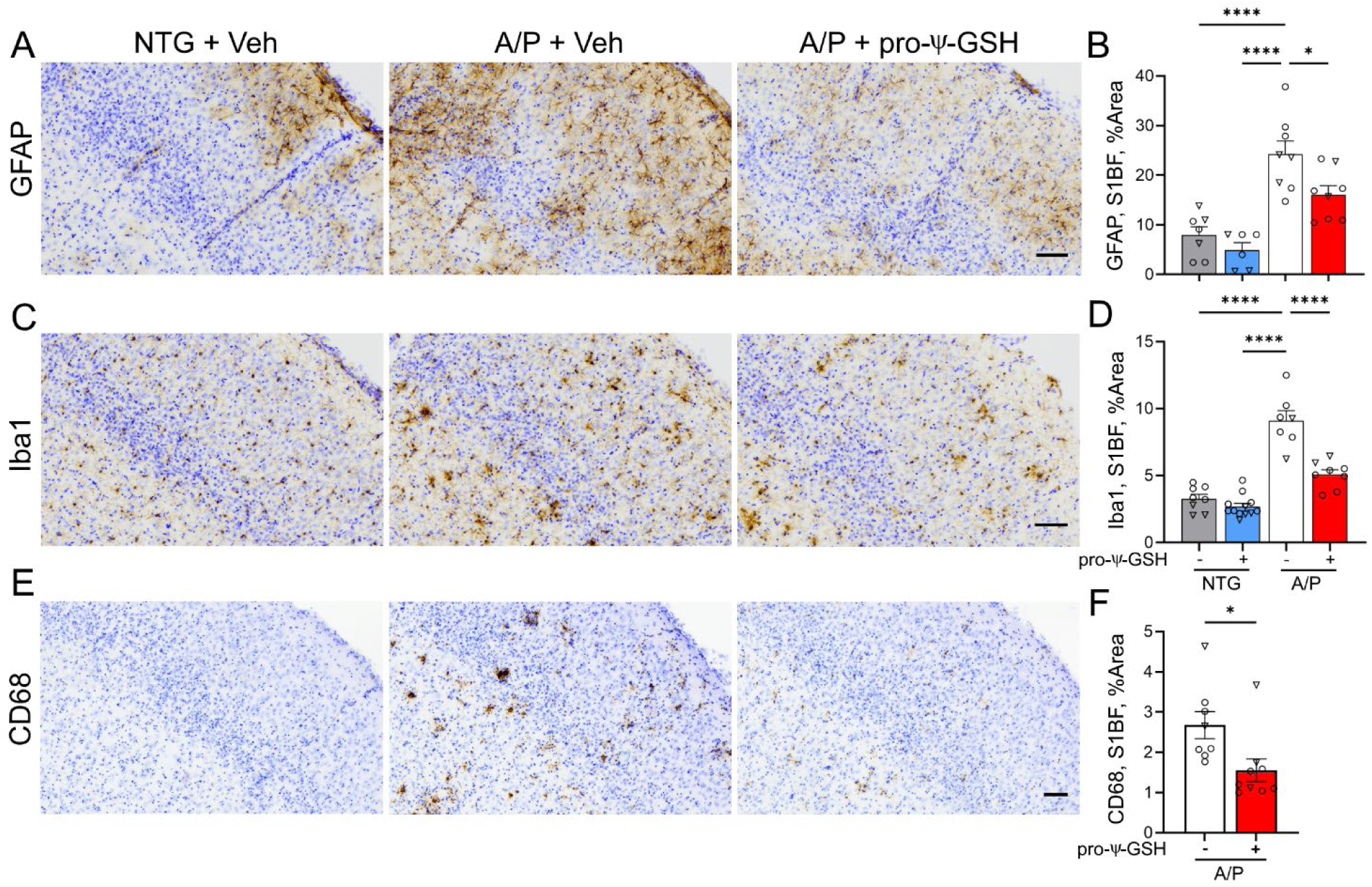
Oral pro-ψ-GSH treatment reduces cortical reactive astrocytosis and microglial activation in the APP/PS1 mouse model of AD. (A, C, E) Representative images of brain sectioned stained for astrocytes (GFAP), total microglia (Iba1) and activated microglia (CD68) in S1BF of NTG and APPswe/PS1ΔE9 (A/P) mouse cohorts. Animals were treated with either Vehicle (Veh) or pro-ψ-GSH for 3 months prior to tissue collection. Scale bar, 100 µm. (B, D. F) Glial reaction in S1BF was quantified by defining the percent area covered by GFAP (B), Iba1(D), or CD68 (F) immunoreactivity. The results show the expected increase in glial activation in the Veh treated A/P mice compared to NTG mice. Consistent with reduced amyloid pathology, oral pro-ψ-GSH treatment significantly reduced astrocyte and microglial activation. Circles are males and triangles are females. Data are shown as mean ± S.E.M. Statistical comparisons between group were performed by one-way ANOVA with Tukey’s post-hoc multiple comparison test. * p < 0.05, ** p < 0.01, *** p < 0.005, **** p < 0.001.

To selectively analyze activated microglia, the brain sections were also stained with an antibody to CD68, which is a marker highly expressed in activated microglia (Figure 6E, F) ^26^. Consistent with the Iba-1 staining, the 12-month-old APP/PS1 mice exhibited prominent CD68 staining in the cortex and hippocampus, while no CD68 staining was seen in NTG littermates. The pro-ψ-GSH treatment significantly reduced CD68 immunoreactivity in APP/PS1 mice (A/P-veh: A/P-pro-ψ-GSH; 2.675 ± 0.34%, 1.554 ± 0.28%; respectively; p < 0.05). The results of Iba-1 and CD68 immunostaining show that pro-ψ-GSH treatment attenuates microglial activation in the cortical region of APP/PS1 mice. Analysis of the hippocampus also showed a similar trend in the prodrug treated animals (Supplementary Figure S2). Microglial activation was significantly increased in saline-treated APP/PS1 brains compared to non-transgenic animals. Oral pro-ψ-GSH treatment rescued against total microglia (Iba1) and content of activated microglia (CD68) in APP/PS1 mice compared to the vehicle controls.

The release of cytokines from activated microglia in AD is known to shift from protective cytokines to pro-inflammatory cytokines ^27^. We evaluated cytokine levels in the cortical homogenates of saline and pro-ψ-GSH treated APP/PS1 mice in relation to non-transgenic controls by multiplex cytokine protein array. The measurement of a comprehensive array of cytokines and chemokines is crucial for understanding of the complex interplay of these molecules in various aspects of disease pathology. Specifically, in AD, measuring cytokines and chemokines is crucial due to their role in neuroinflammation, which influences amyloid precursor protein processing, amyloid aggregation, and tau phosphorylation, thus contributing to the neuropathological features of the disease.^28^ After densitometry and quantification of the homogenates, our results indicated that out of 62 detected cytokines, 29 cytokines were significantly upregulated in saline-treated APP/PS1 mice compared to non-transgenic control littermates (Supplementary Figure S4, and Table S1). In the AD brain, both pro-inflammatory cytokines (IL-2, IL-3, IL-3Rβ, IL-9, IL-13) and chemotactic cytokines (chemokines) such as CXCL13, CXCL16, CX3CL1, CXCL1, MIP-1α/γ, CCL19, CCL17, CCL25 showed increased expression. Additionally, vascular inflammatory markers such as vascular endothelial growth factor (VEGF) and vascular cell adhesion protein 1 (VCAM1) were significantly higher in the APP/PS1-saline group, indicating cerebrovascular inflammation and reduced cerebral blood flow.

Attenuation of cytokine/chemokine response to AD pathology was observed in response to oral pro-ψ-GSH treatment. Specifically, the levels of IL-2, IL-3, IL-6, IL-13, CXC, and CC-chemokines were significantly reduced by the prodrug treatment (Figure 7A, B), correlating with the biochemical benefits offered by the treatment. TNF-α plays a multifunctional role in the inflammatory cascade and two TNF-α receptors are identified that activate different downstream pathways.^29^ Expression of these receptors was reduced by pro-ψ-GSH treatment. Furthermore, vascular abnormalities caused by endothelial inflammation were significantly mitigated by the treatment as evident from the reduction in VCAM1 and VEGF.^30, 31^ It was also interesting to note that the protein expression of insulin-like growth factor binding protein 6 (IGF-BP-6) was affected by the treatment, given the role of Glo-1 in detoxification of oxidative sugar metabolites. Noticeably, activation of M-CSF, a macrophage colony-stimulating factor, was significantly attenuated by pro-ψ-GSH. This growth factor is involved in modulation of AD-related inflammatory processes, and has been shown to offer pro- or anti-inflammatory phenotype depending on the stage of the disease.^32^

**Figure 7.**
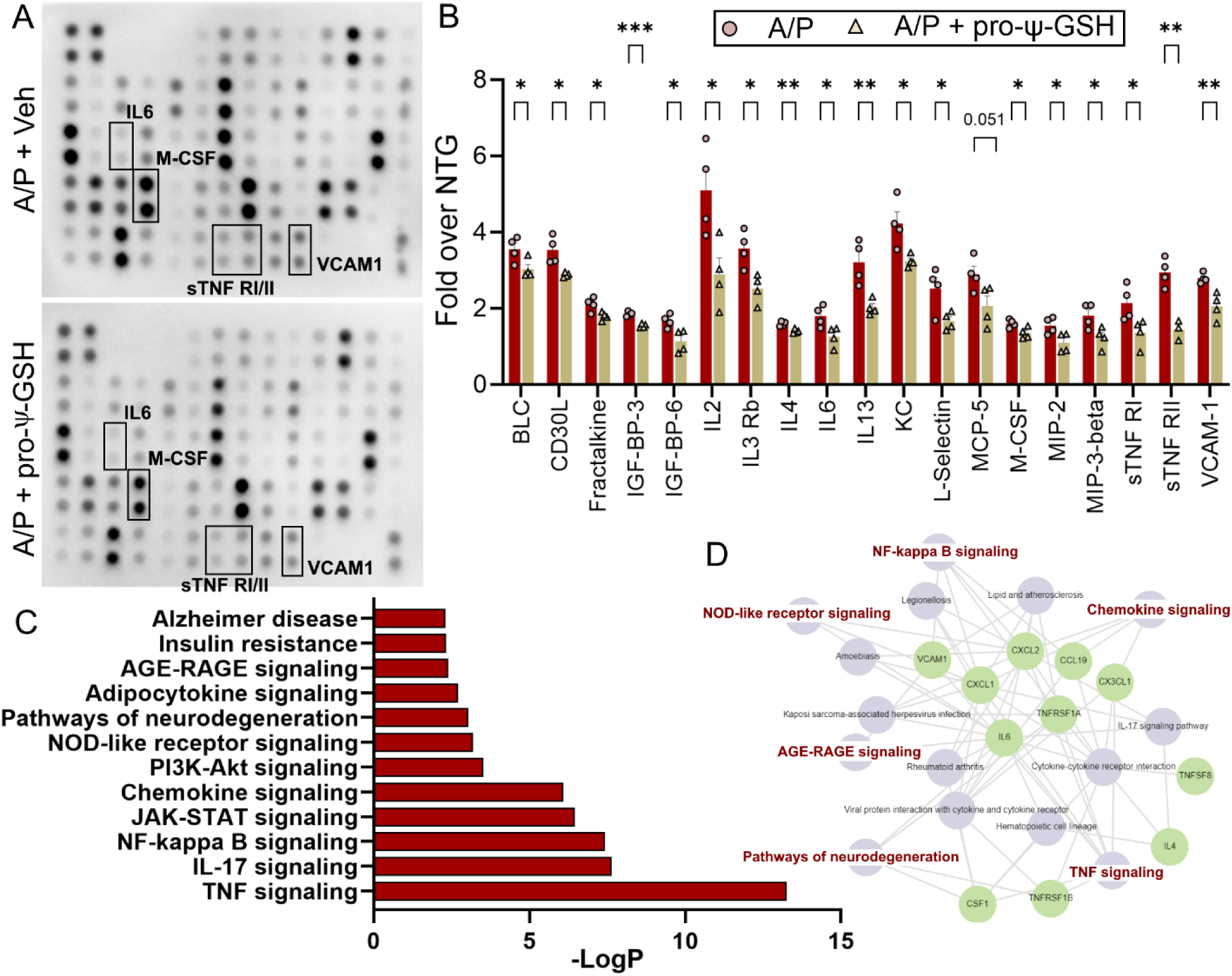
Cytokine array analysis of mice based on genotype and treatment. (A) Representative membranes showing the change in the inflammatory cytokines in brain homogenates from APP/PS1 mice treated with either saline or pro-ψ-GSH. The intensities of the spots were quantified densitometrically by ImageJ and compared to the density of the internal standards. (B) The data is normalized with respect to NTG saline controls. Cytokine analysis showed a significant increase in pro-inflammatory cytokines (IL-1β, IL5, IFN-γ, MIP-α, TNF-α, VCAM-1) in saline treated APP/PS1 mice compared to age-matched NTG controls. Treatment with pro-ψ-GSH reversed the effect of AD pathology in aged APP/PS1 mice, and offered cytokine profile similar to the NTG controls. Data are expressed as mean ± SEM. (C) Pathway analysis using Enrichr online tool displayed modulation of signaling pathways involved in neurodegeneration, specifically Alzheimer’s processes. (D) Display of pathways impacted by pro-ψ-GSH and the involved cytokines. Data are shown as mean ± S.E.M and is representative of two independent experiments. Statistical significance in (B) was determined by two-way ANOVA analysis using Fisher’s LSD multiple comparison or Tukey’s post-hoc test. (* p < 0.05, ** p < 0.01, *** p < 0.001).

The cytokine signature offered by pro-ψ-GSH treatment was analyzed using the Enrichr-KG online analysis tool^33^ (Figure 7C, D). To increase the human relevance of these findings, we analyzed our results using the KEGG human library. We found a positive correlation between cytokine expression level changes to pathways related to TNFα, NFκB, chemokine, and IL-17 signaling. Similarly, when arranged by significance levels, the cytokine changes correlated with insulin resistance and AGE-RAGE signaling pathways. This substantiates the mechanism of neuroprotective action of pro-ψ-GSH. Importantly, the collective modulation of IL6, M-CSF, and sTNF receptors I and II, along with VACM1 by pro-ψ-GSH suggests significant involvement of pathways of neurodegeneration, specifically in AD.

### Treatment with pro-ψ-GSH protects against progressive noradrenergic neurodegeneration in APP/PS1 mice

Our results show that oral pro-ψ-GSH attenuates AD-like biochemical and neuroinflammatory changes in the APP/PS1 model. However, AD is also associated with progressive neurodegeneration that better correlates with the onset and progression of dementia.^34^ While the APP/PS1 model does not exhibit loss of forebrain neurons, this model recapitulates the progressive loss of subcortical monoaminergic neurons seen in human AD.^14, 35^ Thus, we evaluated the integrity of the TH+ noradrenergic (NAergic) neurotransmitter system arising from locus coeruleus (LC) in the APP/PS1 model as a function of oral pro-ψ-GSH treatment.

We analyzed the integrity of NAergic afferents in the cortex by determining the density of TH+ axons in the S1BF via stereological analysis of TH-immunostained brain sections (Figure 8A, B). Representative TH-immunostained sections (Figure 8A) and the quantitative analysis of TH+ afferent density (Figure 8B) show significant loss of TH+ afferents in saline-treated APP/PS1 group (NTG-veh: A/P-veh; 6.263 ± 0.21 × 10^-3^, 2.613 ± 0.22 × 10^-3^ μm/μm^3^; respectively). However, oral pro-ψ-GSH treatment leads to significant protection of TH+ afferents in the APP/PS1 mice (A/P-pro-ψ-GSH; 4.267 ± 0.23 × 10^-3^ μm/μm^3^).

**Figure 8.**
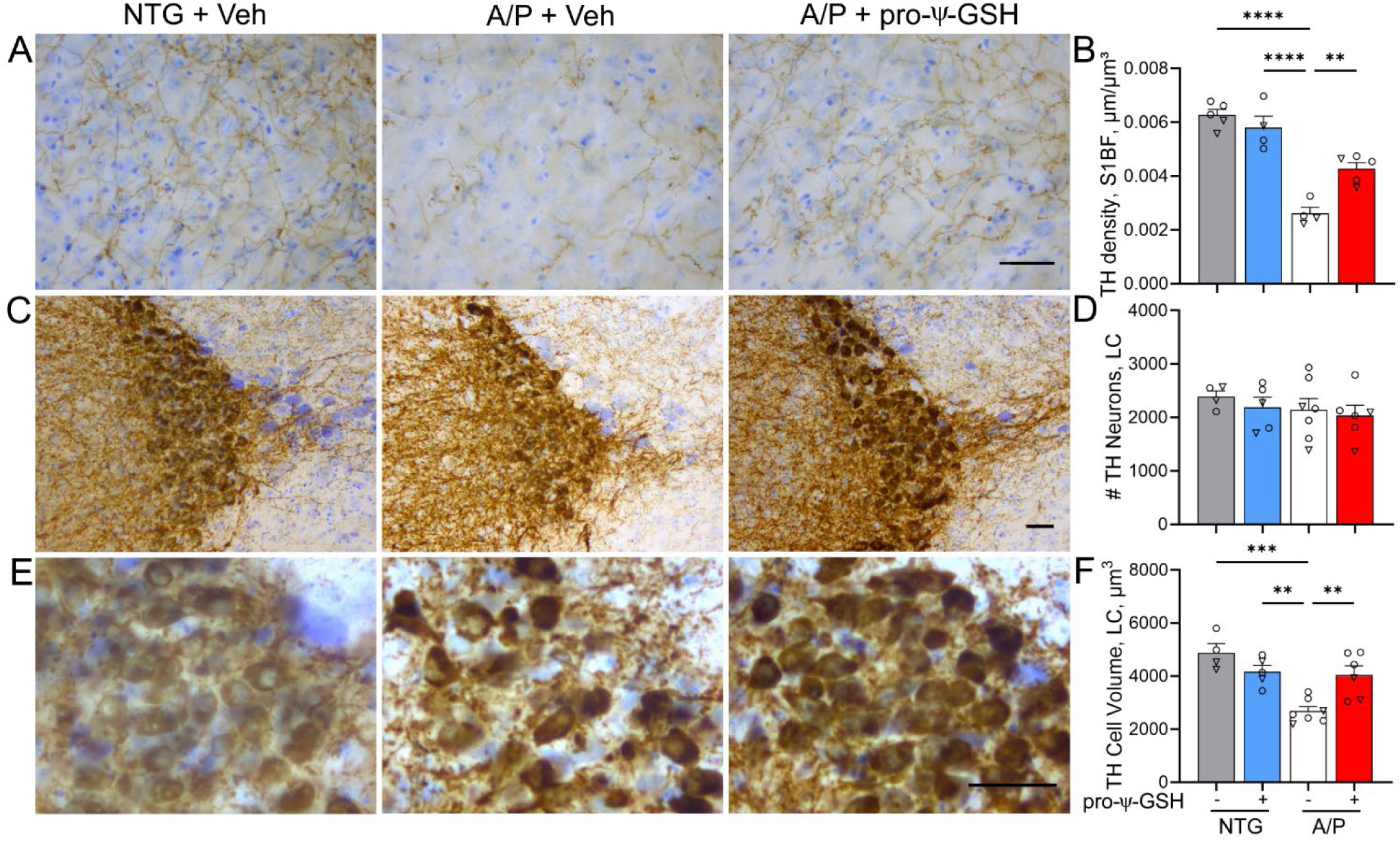
Oral pro-ψ-GSH treatment attenuates the loss of cortical TH+ afferents and atrophy of TH+ neurons in Locus Coeruleus (LC) in the APP/PS1 model. (A) Representative images of TH+ afferents in S1BF of Vehicle (Veh) and pro-ψ-GSH-treated APPswe/PS1ΔE9 (A/P) subjects. Also shown is Veh treated NTG mice. Scale bar, 50 μm. (B) Quantitative analysis of TH+ afferent density show that the TH+ afferent density is reduced in Veh treated A/P animals compared to NTG subjects. Oral pro-ψ-GSH treatment of A/P mice leads to a significant increase in TH+ afferent density compared to Veh treated A/P subjects. (C, E) Representative low (C) and high (E) magnification images of TH+ neurons of the LC. (D) Consistent with prior studies, stereological counting of TH+ neurons in the LC do not show significant neuronal loss in A/P mice. (F) Analysis of TH+ neuron volume in LC show that, consistent with the loss of TH+ afferents in A/P + Veh mice, TH+ neurons are significantly smaller compared to NTG mice. Further, oral pro-ψ-GSH treatment of A/P mice show significant attenuation of neuronal atrophy. Circles are males and triangles are females. Data are mean ± S.E.M. One-way ANOVA with Tukey’s post-hoc test. **p < 0.01, *** p < 0.005, **** p < 0.001.

A previous study^35^ showed a progressive loss of TH+ neurons in the LC occurs after 12 months of age. Thus, analysis of neuronal integrity in LC of the APP/PS1 mice in this study show that all groups exhibit a similar number of TH+ neurons in LC (Figure 8C, D). Similarly, the number of non-TH+ neurons in LC was unchanged across genotypes and treatment groups (Supplementary Figure S5). Despite the lack of neuronal loss in LC, the degeneration of distal NAergic afferents (Figure 8A, B) is predicted to be reflected in neuronal atrophy ^14, 35^. Analysis of neuronal volumes of TH+ neurons confirms that the loss of cortical TH+ afferents in APP/PS1 mice is associated with a significant reduction in cell volume, indicating significant neuronal atrophy (Figure 8E, F). Consistent with the neuroprotective effects of pro-ψ-GSH, neuronal volumes in pro-ψ-GSH treated APP/PS1 mice were comparable to the non-transgenic controls (NTG-veh: A /P-veh: A/P-pro-ψ-GSH; 4893 ± 340, 2680 ± 169, 4048 ± 340 μm^3^, respectively). Treatment of non-transgenic mice with pro-ψ-GSH did not have any significant effect on TH+ fiber density, cell number, or volume when compared to the corresponding vehicle controls (Figure 8 and Supplementary Figure S6).

### Pro-ψ-GSH normalized metabolite changes caused by AD pathology in APP/PS1 mouse brain tissue

In addition to oxidative stress, neuroinflammation, and neurodegeneration, metabolite alterations are one of the earliest signs of the disease.^36, 37^ In this study, we examined metabolite profiles of the hippocampal region of our animal cohort based on the genotype and treatment (NTG-saline; A/P-saline; A/P-pro-ψ-GSH and NTG-pro-ψ-GSH) to identify metabolic pathways associated with disease pathology, GSH supplementation, and Glo-1 pathway modulation. The schematic of this study using the commercial Biocrates kit is shown in Figure 9A. Metabolic profiles of the hippocampal region from the treatment groups were first aligned together to perform sample classification by multivariate data analysis. An initial principal component analysis (PCA) was performed to check the quality of the analysis, identify outliers, and ensure proper grouping of the QC samples. Partial least squares discriminant analysis (PLS-DA) of the data sets offered an improved separation of different study groups (Figure 9B). This model yielded satisfactory values for the quality parameters, goodness of fit R^2^ as 91%, and accuracy of 60% using 5-fold cross-validation and two components, and yielded metabolite markers that significantly contributed to the differentiation between the treatment groups. We identified over 350 metabolites from 26 biochemical classes. Significant alterations in metabolites such as phospholipids, fatty acids, purine metabolites, and amino acids were detected, as previously reported for this model.^38, 39^ Increased levels of ceramides (Cer) and sphingomyelins (SM) found in APP/PS1 brains were reduced by the prodrug treatment, while levels of polyunsaturated fatty acid (PUFA), important for brain function, were increased (Figure 9C). Discriminant metabolites were then selected according to the VIP value (VIP score > 1.5) for each PLS-DA model.

**Figure 9.**
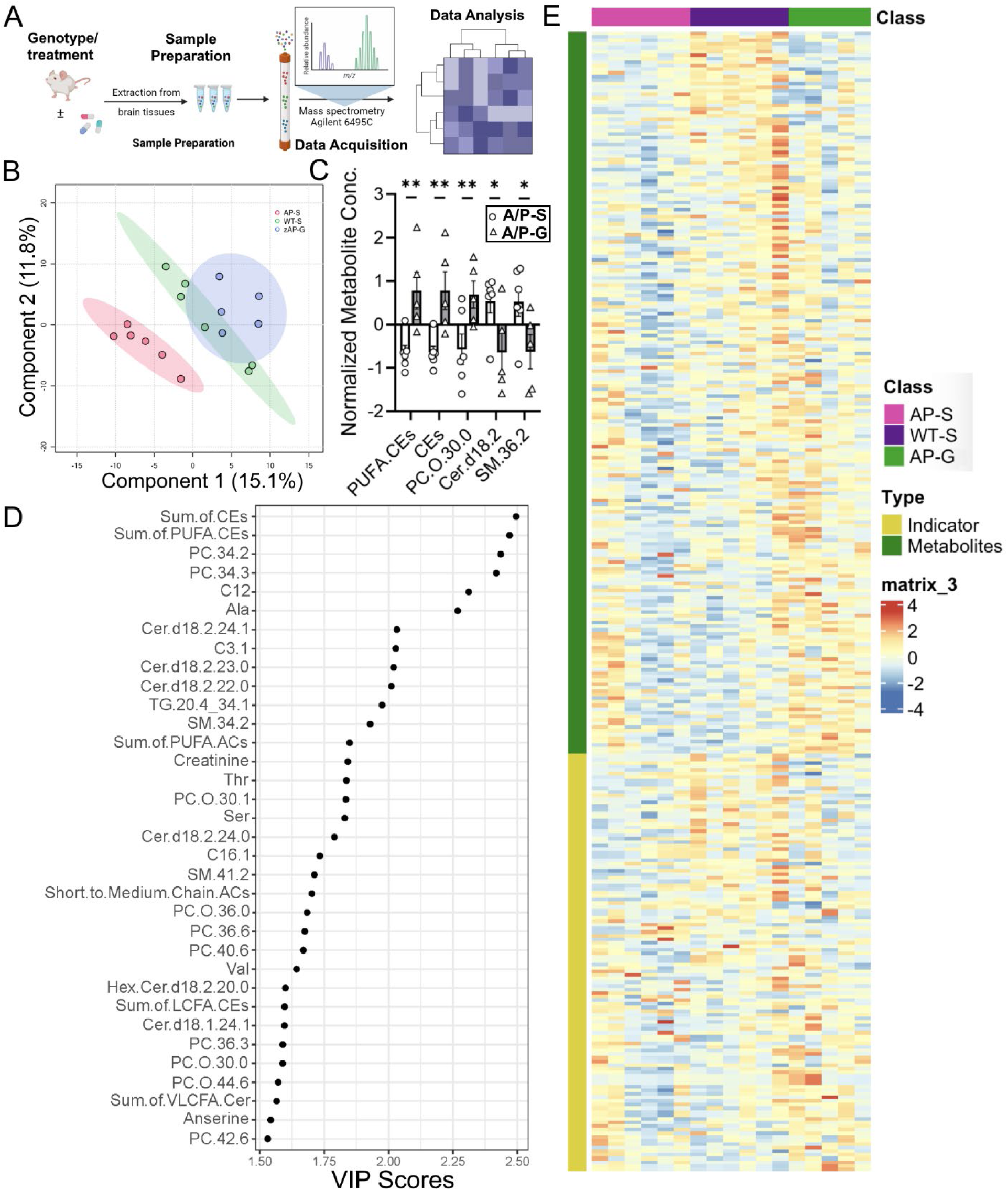
Metabolite alterations caused by AD pathology in APP/PS1 mice were reversed by pro-ψ-GSH treatment. (A) Schematic of the metabolomics study. (B) PLS-DA plot classifying the hippocampal metabolites from NTG and APP/PS1 (AP) mice treated with pro-ψ-GSH (AP and AP/G, resp.). (C) Bar graph showing top metabolites that are affected by the prodrug treatment in APP/PS1 (A/P-G) compared to corresponding vehicle treated mice (A/P-S) with established role in AD pathology. PUFA-CEs = polyunsaturated fatty acid cholesteryl esters, CEs = sum of cholesteryl esters, PC.O = phosphatidylcholine, cer = ceramide, SM = sphigomyelins. (D) Metabolite profiling analysis by VIP scores (> 1.5) provided top metabolites contributing significantly to variations within groups. The relative contribution of the metabolites is proportional to the VIP score. (E) Heat maps showing changes in top 50 metabolites in NTG and APP/PS1 groups treated with either saline or pro-ψ-GSH. Each column represents an individual animal within the group. Hierarchical clustering found NTG-saline group closer to the A/P + pro-ψ-GSH group, suggesting beneficial effect of the treatment. Data are represented as the mean ± SEM. Statistical significance was derived by Student t-test, * p < 0.05, ** p < 0.01, n = 5-6.

Amongst the metabolites of significant importance were metaboindicators — sum of PUFA-cholesteryl esters and cholesteryl esters. Although alterations in various classes of lipids have been detected in AD brains, changes in unsaturated fatty acids have been associated with AD progression.^40^ The effect of pro-ψ-GSH treatment on the levels of such lipids, which are involved in the regulation of various biological processes, is thus noteworthy. Further, some of the metabolites suggest treatment-specific alterations in glycosphingolipids such as glycosylated ceramide (hexosylcer-amide, Hex-Cer) that are known to contribute toward oxido-inflammatory stress in diabetes and neurodegeneration.^41, 42^ Multi-variate analysis using the Receiver Operating Characteristic (ROC) curve provided a biomarker model using a sum of PUFA-CEs and the sum of CEs offered an AUC of ∼1 and thus, demonstrates the potential of metabolomics screening for understanding treatment-specific biomarkers (Supplementary Figure S7). There are additional metabolites related to triglycerides, sphingomyelins, and ceramide that also offer good statistical correlation with the treatment effect (Supplementary Table S2). This is further substantiated by fold-change analysis caused by the treatment with respect to vehicle-treated APP/PS1 mice. As expected, supplementation with pro-ψ-GSH improved the synthesis of cysteine and other amino acids, purines, and carnosine. The treatment reduced the levels of saturated fatty acids, deemed beneficial for preventing Alzheimer’s progression.

## DISCUSSION

Glycation-induced oxidative stress is a major contributor to the neurodegeneration observed in AD. The loss of reduced GSH and compromised Glo-1 function contribute to increased oxidative stress found in the AD brain. We show that these changes are recapitulated in the APP/PS1 mouse model of AD. While intraperitoneal ψ-GSH reverses cognitive dysfunction in APP/PS1 mice when initiated at pre-symptomatic and post-symptomatic stages,^16^ the compound did not improve cognition when administered orally, requiring synthesis of orally available prodrugs of ψ-GSH.^17^ We now show the effect of oral ψ-GSH prodrug (pro-ψ-GSH) on AD-like neuropathology, oxidative stress, inflammation, and cognitive impairment in the APP/PS1 mouse model when initiated after the establishment of progressive pathology.

APP/PS1 mice overexpress human transgenes for APP and PSEN1, increasing deposition of amyloid plaques with age, which leads to impaired spatial and working memory at around seven months of age.^43^ A combined battery of cognitive tests showed cognitive deficits in saline treated mice transgenic mice compared to nontransgenic mice at the same age (12 months). Evaluation of pro-ψ-GSH treated mice established beneficial effect of the treatment on hippocampal and cortical activity dependent cognitive behavior. Immunohistochemical and ELISA analysis corroborated with improvement in cognitive performance, and clearly demonstrated attenuation of Aβ pathology by oral pro-ψ-GSH, even when initiated following the onset of amyloid pathology.

Glo-1 activity is crucial in mitigating oxidative stress induced by reactive dicarbonyls such as MG and proteolytically resistant AGEs.^44, 45^ A bioavailable GSH analog such as pro-ψ-GSH is expected to alleviate carbonyl stress, partly by restoration of compromised Glo-1 function. Thus, we show that treatment with oral pro-ψ-GSH, not only improved brain GSH levels, but also restored Glo-1 activity and reduced accumulation of MG/AGEs in APP/PS1 mouse brain. Pro-ψ-GSH treatment also reduced other indices of oxidative stress such as lipid and protein oxidation products, and augmented GSH-dependent glutathione peroxidase enzyme. It is thus apparent that supplementation with a GSH analog could have effects beyond the Glo-1 pathway and thus could have multiple benefits due to restoration of GSH-dependent enzymatic functions.

The increase in glycated intermediates resulting from reduced Glo-1 activity is linked to inflammatory response via activation of the receptor for AGEs,^46^ eventually contributing to neuronal dysfunction and neurodegeneration found in AD. Astrocytes and microglia play an important role in maintaining brain homeostasis and undergo activation in response to neurodegenerative processes. Our neuropathological analysis confirmed the anti-inflammatory activity of oral pro-ψ-GSH from reduction in the levels of activated astrocytes and microglia, even at this late stage in the symptomatic APP/PS1 mouse model. Further analysis of neuroinflammatory cytokines captured a major inflammatory storm induced by the disease pathology. Increased levels of pro-inflammatory cytokines and chemokines such as IL-2, IL-6, CXCL16, and vascular VCAM1 were found in saline-treated APP/PS1 brains compared to the age-matched controls. The results corroborated with extensive research in AD mouse models that have identified similar expression changes in cytokines and chemokines, playing pro- or anti-inflammatory roles depending on the disease stage and presence of disease risk factors.^27^ Thus, this transgenic mouse model accurately replicates some of the pathological inflammatory features seen in the AD patient brain. Pro-ψ-GSH treatment significantly attenuated the inflammatory cytokine levels to levels found in non-transgenic animals. Interestingly, the pro-ψ-GSH treatment reduced the levels of M-CSF, a macrophage stimulating factor, which has been pursued as a putative biomarker for mild cognitive impairment,^47^ demonstrating utility of this approach to identify treatment-specific biomarkers. Further, the inhibitory IGF-BP-6 proteins for insulin-like growth factor 1 accumulate in hippocampal pyramidal neurons and in amyloid plaques in the AD brain and are shown to be responsible for early cognitive deficits.^48^ Methylglyoxal is one of the toxic metabolites of sugar oxidation and clearance of such neurotoxic intermediate and AGEs with Glo-1 supplementation could thus provide an effective strategy for improving insulin signaling in the AD brain.

Restoration of the Glo-1 pathway is expected to affect the activation of RAGE-induced inflammatory signaling. RAGE is a multi-ligand receptor belonging to the immunoglobulin superfamily, whose engagement leads to sustained cellular dysfunction and tissue damage. Involvement of RAGE further contributes to vicious oxidative stress cycle and contributes to neuroinflammation by nuclear factor-kB (NF-kB) up-regulation.^49^ This ultimately leads to transcription and release of cytokines such as TNF-alpha and chemokines, inducing a prolonged activation of cell damage signaling mechanisms.^50^ The biochemical analysis of the brain tissues corroborated with the predicted enrichment of RAGE-associated neurodegenerative pathways in saline treated APP/PS1 mice, which was effectively normalized by pro-ψ-GSH. Such mitigation of inflammatory stress by the prodrug contributed to protection against progressive monoaminergic neuronal loss, even when initiated after the onset of neurodegeneration.

Metabolic changes are evident at early stages of AD by changes in glucose utilization detected using fluorodeoxyglucose positron emission tomography (FDG-PET) in patients suffering from mild cognitive impairment. Additionally, perturbations in other metabolic networks such as neurotransmitter synthesis, amino acid metabolism, tricarboxylic acid (TCA) cycle, and fatty acid/lipid metabolism are reported in MCI patients and transgenic AD animal models.^51, 52^ Dysregulation of brain metabolism is considered a risk factor for AD, which is further complicated by disease heterogeneity due to genetic diversity, environmental and lifestyle factors in humans. Transgenic AD animal models show significant overlap in the affected metabolic pathways identified in AD patients and provide a critical understanding of the disease progression.^30, 38^ Our metabolomics study confirmed the replication of metabolic dysregulation found in human AD brain in this APP/PS1 model and showed reversal of metabolic alterations by the prodrug treatment. The reported changes in brain lipid profile, ceramide, amino acids were reproduced in this analysis and were effectively normalized by oral pro-ψ-GSH. The analysis has provided key metabolites such as sphigomyelins and cholesteryl esters, and associated pathways that could potentially serve as treatment-specific markers in a clinical setting for Glo-1/AGE targeted therapeutics. Additional investigations are however needed to validate these markers based on the disease stage and various brain regions. Collectively, the data provides insights into potential metabolic pathways affected by the prodrug treatment.

## CONCLUSION

In this study, we demonstrate that oral pro-ψ-GSH treatment attenuates AD-related abnormalities in the APP/PS1 model even when treatment is started after overt amyloid pathology. APP/PS1 mice display increased oxidative stress, reduced GSH levels, and impaired Glo-1 function. Along with the increased amyloid pathology and neuroinflammation, these mice also develop cognitive deficits and progressive neurodegeneration of MAergic neurotransmitter systems.^14, 35, 53^ Orally available prodrug of the Glo-1 substrate ψ-GSH, ameliorated cognitive behavioral, neuropathological, and biochemical changes apparent in these transgenic mice. Pro-ψ-GSH offers pharmacokinetic advantage over the parent compound, ψ-GSH, in brain delivery of the bioactive species, however the brain concentration of the intact prodrug was marginally elevated, partly because of its rapid conversion to ψ-GSH either in the periphery or after delivery to the brain. Future design of brain-targeted analogs of pro-ψ-GSH could lend additional benefit to this therapeutic approach. The results of this study provide support for additional investigations directed toward development of ψ-GSH-based orally bioavailable therapeutics as potential therapy for AD. Based on our recent investigations in Glo-1 knockout mice, such therapeutics could have pharmacological targets beyond Glo-1, requiring rigorous mechanistic explorations. Given the multifactorial nature of GSH-based therapeutics, additional efficacy experimentation against multiple aspects of AD pathology, such as in the presence of tauopathy, are highly warranted. Such therapeutic approach is expected to provide much needed disease modification to affect disease course in the clinic.

## MATERIALS AND METHODS

### Synthesis of pro-ψ-GSH

Pro-ψ-GSH was synthesized in our laboratory using a previously described synthetic method.^17^ Authentication of the compound was through NMR analysis and purity (> 96% pure) of pro-ψ-GSH was confirmed by reverse phase HPLC analysis.

### Mice

APP/PS1 transgenic mice [B6.Cg-Tg(APPswe,PSEN1dE9)85Dbo/Mmjax; 034829-JAX] breeders were obtained from Jackson Laboratory and maintained at the University of Minnesota. All experimental procedures and animal handling were performed in accordance with the national ethics guidelines and complied with the protocol requirements of the Institutional Animal Care and Use Committee (IACUC) of the University of Minnesota, Minneapolis, MN. The animals were housed in four per cage in the university animal facility under controlled environmental conditions on a 12 h-12 h light-dark cycle and were allowed access to food and water ad libitum. APP/PS1 transgenic and age-matched non-transgenic mice were used in this study. The mice were treated with either saline or pro-ψ-GSH via oral gavage (250 mg/kg in sterile saline) three times a week for three months starting at 9 months of age (N = 12-14 per treatment group). The choice of the dose was based on the previous pharmacokinetic evaluation and efficacy evaluation of the prodrug.^17^ After completion of the treatment course and behavioral analysis at 12 months of age, the cohort of mice was sacrificed under isoflurane anesthesia, and brain tissues were collected for biochemistry and neuropathological analysis.^14, 54^

### Behavior Studies

#### Y-maze

**Spontaneous alternation test** was performed to assess short-term spatial working memory on the Y-maze. The maze used in the study was constructed of acrylic plastic and measured 30L × 10W × 20H cm for each of its three arms, which converged at an equal angle. Visual cues were placed on the maze walls. Prior to the test, the mice were individually placed at the end of one arm of the maze facing the wall and their movements were recorded for eight minutes. The number of times the mice visited each arm was recorded, and alternation was defined as the number of times the mouse visited each arm sequentially, without repeating any arm. Spontaneous alternation was calculated as the percentage of total arm visits minus two, with a smaller percentage indicating memory impairment. The equation used is as follows: % spontaneous alternation = ((# spontaneous alternations) / (Total arm entries - 2)) * 100.

#### Y-maze

**Spatial recognition test** for spatial recognition memory was also performed on the Y-maze. To perform this test, one arm of the Y-shaped maze was obstructed, which enabled the animal to explore the remaining two arms for 10 minutes. The animal’s memory function was subsequently tested 24 hours later by returning it to the maze with all arms open and observing its inclination to spend time in new or known arms for 5 minutes. This test serves as an effective tool for assessing memory function in mice.

### Novel Object Recognition Test

The Novel Object Recognition (NOR) test in mice is a behavioral assay used to evaluate recognition memory. The test typically involves three phases: habituation, training, and testing. During habituation, mice were allowed to explore an empty arena to become familiar with the environment. The training phase (5 min) follows, where mice were presented with two identical objects and allowed to explore. In the testing phase (5 min), one of the familiar objects was replaced with a new object, with a different color and shape, and the time spent exploring the novel object was recorded as an index of recognition memory. The test session was carried out 1 h after training. The percent time spent with the novel object was calculated to analyze the interaction of the animal with the new object compared to the familiar object.

### Barnes maze test

The Barnes Maze behavioral test is widely used to assess spatial learning and memory in rodents, particularly in mice. The protocol involved placing the animal on a circular platform with multiple holes around the perimeter, one of which led to an escape box. The maze consisted of an elevated circular platform (90 cm above ground and 100 cm in diameter) with 20 equally spaced holes (diameter = 5 cm) situated around its perimeter. The maze surface was white. An escape box was hidden beneath one of the holes and four distinctive extra-maze reference cues were placed around the maze, one on each side (large black letters on a white background). The testing took place in a brightly lit room with a 300-watt light directly overhead to encourage the mice to prefer dark, enclosed spaces to bright, open areas. Animals are motivated to find this escape hole to avoid aversive stimuli, such as bright lights.^22, 55^ All animals began each trial positioned at the center of the maze. To prevent the mice from following scent trails, the maze was cleaned with a 70% ethanol solution between trials. Briefly, on the first day of testing, all mice were administered two habituation trials. Mice were placed in a start chamber located at the center of the maze for 10 s. Subsequently, the start box was lifted, and the mice were gently guided to the escape box, where they were allowed to enter on their own. After entering the escape box, the mice remained there for 30 s before being returned to their home cage for a 2-minute inter-trial interval. From days 2 to 4, the mice underwent the acquisition phase, which consisted of four trials per day changing the orientation of the maze, and escape box, with a 15-minute inter-trial interval between trials. The trials ended when the animal entered the escape box or when a maximum trial length of 3 min was reached. Mice were allowed 30 s in the escape box. If the animal did not find the escape box within the allotted 3-minute trial period, they were gently guided to the escape box and allowed 30 s inside. On day 5, the animals underwent one probe trial, the escape box was removed from the target hole that had been previously learned, and the mice were given 90 s to explore the maze. The percentage of time spent in the correct quadrant during the first 60 s of the probe trial was then calculated.

### Homogenization

Sections of the cortex and hippocampus were excised from the brains. Half of the cortex was utilized for western blots, which was homogenized in RIPA buffer, while the other half was homogenized in PBS buffer for biochemical experiments. PBS buffer was used to homogenize the hippocampal areas for biochemical tests. In brief, PBS or RIPA buffer was added to the brain tissue and then homogenized using a mechanical homogenizer at 6,000 rpm speed. Solution was then centrifuged at 15,000 rpm for 20 min at 4° C and the clear supernatant was collected and stored at -80° C.

### Western Blotting

RIPA buffer containing protease inhibitor (Roche) was used to lyse the cortex. Protein concentrations were measured using the BCA Protein Assay Kit (Thermo Fisher Scientific). After the proteins were denatured, the same quantity of protein (40 µg/lane) from each group shown in the images was separated by SDS-PAGE and then transferred using a wet transfer method to a polyvinylidene fluoride (PVDF) membrane for immunoblot studies. After applying 5% milk to block the PVDF membrane for two hours, antibodies against Glo-1 (ab137098), Glo-2 (AF5944), and α-tubulin (ab4074) were incubated at 4 °C overnight. ECL substrate and an appropriate secondary antibody (1:5000) linked to horseradish peroxidase were used to label the primary antibodies.

### AGE ELISA assay

AGE levels in brain homogenates were determined using a commercial ELISA kit (XpressBio, Cat. No. XPEM0716, Frederick, MD, USA). To do this, 100 µL of the homogenate (diluted 10-fold with dilution buffer) was added to a 96-well plate pre-coated with the anti-AGE antibody and incubated at 37 °C for 90 min. The plate was then washed, and 100 µL of biotin-labeled anti-AGE antibody was added to each well. After another round of washing, 100 µL of HRP-conjugated streptavidin was added and incubated for 30 minutes at 37 °C. The plate was then washed again and incubated with TMB substrate for 20 min before quenching the reaction with the provided solution. The absorbance was measured at 450 nm, which indicated the AGE content.

### Amyloid beta ELISA assay

Sandwich ELISA was used to quantify the levels of soluble (PBS fractions) and insoluble (guanidine fractions) Aβ in the cortical region according to the manufacturer’s instructions (Invitrogen-KHB3441).^14, 54^ For quantification of insoluble Aβ levels in the tissue homogenates, the cell pellet obtained after PBS homogenization was dissolved in 5 M GnCl-50 mM Tris HCl at pH 8.0 and was allowed to sit at room temperature (RT) for 15 min with vortexing in between, then centrifuged at 15,000 rpm for 15 min at 4° C and supernatant was stored at -80° C.

Briefly, to a 96-well plate, 50 μL of either guanidine or PBS fractions of brain homogenates were added containing an Aβ detection antibody for three hours at room temperature. Subsequently, 100 μL of anti-rabbit IgG HRP was incubated with the short interim washing procedures. Using a SpectraMax M5e microplate reader, absorbance was measured at 450 nm following the addition of 100 μL of chromogenic solution to each well. The protein content of each sample was used to normalize the levels of Aβ_1-42_.

### Quantification of GSH

Quantification of reduced and oxidized GSH levels in the cortex of mice was performed using a GSH assay kit (Cayman-703002).^14^ To determine the amount of total and oxidized GSH, 10 μL of brain tissue homogenates lysed in PBS were first subjected to equal volumes of metaphosphoric acid (MPA) reagent for deproteinization of the samples. The deproteinated samples were then used in an enzymatic GSH recycling assay using glutathione reductase. This method of determining the GSH level involves derivatization of GSH thiol by 2-nitro-5-thiobenzoic acid (DTNB), resulting in the formation of yellow colored 5-thio-2-nitrobenzoic acid (TNB), and measurement of the absorbance at 410 nm. The data was normalized to the respective protein concentration in each sample.

### Detection of Glutathione Peroxidase (GPx) Activity

A commercial assay kit was used to determine the activity of GPx (Cayman-703102) as per the manufacturer’s instructions. In brief, 20 μL of brain homogenates were incubated with 50 μL of assay buffer, 50 μL of co-substrate mixture, and 50 μL of NADPH. The reaction was quickly initiated by addition of cumene hydroperoxide to all the wells and the absorbance was read at 340 nm using a SpectraMax M5e microplate reader. The absorbance values were normalized to the protein concentrations of the samples to determine enzymatic activity.

### Detection of Methylglyoxal levels using LC-MS/MS

Methylglyoxal content was determined by LC-MS/MS analysis using 1,2-diaminobenzene (DB) as a derivatizing agent, as described previously.^14^ Briefly, the samples were modified by adding 20 µL ice-cold TCA-saline, 40 µL water, and 10 µL 400 nM 2,3-hexanedione (internal standard) to 40 μL mouse brain homogenates. The samples were then centrifuged at 10,000× g for 10 min at 4 °C and the supernatants were analyzed using LC-MS/MS. The LC-MS/MS conditions were as follows: Agilent 1260 HPLC device coupled with an AB Sciex QTRAP 5500 mass spectrometer, Phenomenex Kinetex C18 column (50 × 2.1 mm, 2.6 µm), 0.1% formic acid in water (mobile phase A), and 0.1% formic acid in acetonitrile (mobile phase B), with a flow rate of 0.5 mL/min. A positive-mode electrospray ionization source was used for analyte detection with previously described mass transitions.

### Glyoxalase Assay

Enzyme activity was quantified using a two-step protocol. To begin, Solution Mixture - I (2.5 mL of 100 mM Sodium phosphate, 0.5 mL of 20 mM GSH, 0.5 mL of 20 mM MG, and 1.4 mL of H2O) was prepared and incubated for 10 minutes at RT. Following incubation, 5 μL of brain tissue homogenate was added to a UV multi-well plate (Corning, Cat. No. 3679), along with 95 μL of Solution Mixture - I. The plate was read at 240 nm every 10 s for 5 min. After 5 min, the plate was incubated at 37 °C for additional 10 min, and the final absorbance was read at 240 nm. Subsequently, 5 μL of the sample was added to a 95 μL of Solution Mixture - II (2.5 mL of 100 mM Sodium phosphate, 0.5 mL of 20 mM MG, and 1.4 mL of H2O) and read immediately at 240 nm every 10 seconds for 5 minutes. After 5 min, the plate was incubated at RT for 10 min, and readings were taken. Finally, 2 μL of 100 mM GSH was added to the plate and readings were taken immediately at 240 nm every 10 s for 5 min and at 10 minutes for end-point readings.

### TBARS Assay

Lipid peroxidation levels in the mouse brain were measured using a TBARS assay kit (Cayman-10009055). Briefly, 10 μL of brain homogenate in RIPA buffer was treated for one hour in boiling water with 10 μL of SDS and 400 μL of a color reagent containing thiobarbituric acid. The vials were then removed immediately and left in an ice bath for ten minutes. The supernatants were measured in duplicate at 530 and 550 nm for excitation and emission, respectively. The MDA levels were quantified from a standard curve and normalized with respective protein concentrations for each sample.

### Analysis of Cytokines using Cytokine Array kit

The present study utilized mouse brain tissue lysates from various genotypes and treatments to detect circulating cytokines and chemokines using the Mouse Cytokine Antibody Array C3 kit (RayBiotech, Norcross, GA), following the manufacturer’s instructions. In brief, the membranes coated with capture antibodies were blocked with the provided blocking buffer and then incubated at 4 °C with tissue lysates that were diluted with blocking buffer. Following this, the membranes were washed with the provided washing buffer and incubated with a biotinylated detection antibody cocktail 1:1000 overnight at 4 °C. The next step involved washing the membranes once again and incubating them with streptavidin-HRP 1:5,000 for two hours, followed by the development using detection buffers provided in the kit. The immunoblot images were captured and visualized using a BioRad Molecular Imager GelDoc, and the intensity of each spot was analyzed using ImageJ software (version 1.54a).

### Untargeted Metabolomics

Untargeted metabolites were profiled following a standard protocol of the MxP Quant 500 kit (Biocrates) at the Center for Metabolomics and Proteomics (CMSP) at the University of Minnesota. In total, 630 metabolites covering 14 types of small molecules and 9 distinct lipid classes were analyzed using the MxP® Quant 500 kit (Biocrates) following the manufacturer’s protocol. The MetIDQ software (Biocrates) was used to summarize 12 different lipid classes into 9 by merging lysophosphatidylcholines and phosphatidylcholines into glycerophospholipids, as well as hexosyl-, dihexosyl-, and trihexosylceramides into glycosylceramides. The samples were analyzed on an Agilent 6495C mass spectrometer (Sciex, CA) coupled to an Agilent 1290 Infinity II HPLC with an ESI source. Frozen hippocampal brain samples were homogenized in isopropanol, and derivatized using 5% phenyl isothiocyanate in pyridine solution in the presence of internal standards to derivatize amino acids and biogenic amines. Metabolites were then extracted with ammonium acetate in methanol prior to analysis. The labeled amines were analyzed by LC-MS/MS in positive and negative modes, while the other metabolites were detected by flow injection tandem mass spectrometry in the positive ESI mode. Data was recorded using the Analyst (Sciex) software suite and transferred to the MetIDQ software (version Oxygen-DB110-3005) for further data processing, including technical validation, quantification, and data export. All metabolites were identified using isotopically labeled internal standards and multiple reaction monitoring (MRM) using optimized MS conditions as provided by Biocrates.

Sample outliers were removed from the protein quantification table based on PCA analysis implemented in MetaboAnalyst 6.0 (https://www.metaboanalyst.ca/). The processed profile file was re-imported into MetaboAnalyst 6.0 for data filtering (default setting with unchecked Reliability filter, “IQR” for variance filter, and “Mean intensity value” for Abundance filter), normalization (default setting with “Auto scaling” for data scaling) and downstream statistical analysis, including testing for differential abundance, principle component analysis (PCA) and Partial Lease Square – Discriminant Analysis (PLS-DA) using default parameters. Additional biomarker analysis between APP/PS1-vehicle and APP/PS1-pro-ψ-GSH groups was run using the following setting: Compute and include metabolite ratios – Top 50, Data Scaling – Auto Scaling, and Classical univariate ROC curve analyses. Result visualization was created by R packages MetaboAnalystR (v4.0.0) and ggplot2 (v3.5.1).

### Immunohistochemistry

The brains were processed for immunohistochemistry as previously described.^14, 35^ Briefly, the brains were fixed in 4% paraformaldehyde and cryoprotected in 30% Sucrose/PBS solution. Brains were coronally sectioned (40 μm) on a freezing sliding microtome (Leica). The sections were serially collected on a 12 well plates such at each well contained every 12^th^ section through the entire brain. To detect antigens of interest, the sections were incubated in primary antibodies followed by the ABC method (Vector Laboratories, Burlingame, CA, USA) using the chromogen, 3,30 -diaminobenzidine (DAB; Sigma Aldrich, St. Louis, MO, USA) for visualization. Antigen retrieval was performed using a Rodent DeCloaker (BioLegend, San Diego, CA, USA) for all samples, with an additional 70% formic acid pre-treatment for detecting Aβ plaque markers. Primary antibodies used are: clone 4G8 anti-Aβ17–24 mouse monoclonal antibody (BioLegend), GFAP anti-rabbit polyclonal antibody (Dako, Glostrup Kommune, Denmark), Iba1 anti-rabbit polyclonal antibody (Wako, Japan), CD68 anti-rabbit monoclonal antibody (Abcam, USA) and TH (tyrosine hydroxylase) anti-rabbit polyclonal antibody (Millipore, Burlington, MA, USA). Diluted cresyl violet (CV) was used to counterstain nuclei of neurons.

### Quantitative analysis of amyloid pathology and glial activation

Sections stained for Aβ, GFAP, CD68 and Iba1 were mounted on slides and scanned using Motic EasyScan One (Motic Digital Pathology, Emeryville, CA) slide scanner. The immunoreactivity was quantified using HALO Image Analysis Platform (version 4.0.5107) and Halo AI (version 4.0.5107) (Indica Labs, Inc.). The brain area covered by Aβ deposits (4G8), reactive astrocytes (GFAP), and microglia (Iba1) within the regions of interest (ROI) was measured using the HALO Area Quantification. Image optimization, when used, was applied consistently to all sampled subjects. The ROIs, determined using the Mouse Brain Stereotaxic Coordinates^35^ as the reference, include the barrel field region of primary somatosensory cortex (S1BF; sections between bregma −0.10 to −1.22 mm, posterior to the anterior commissure and anterior to hippocampus) and dorsal hippocampus (dentate, CA1, CA2/3; sections between bregma −1.46 to −2.18 mm). For each animal, we analyzed 4-6 brain sections from a well containing every 12^th^ sections through the entire brain.

### Stereological Analysis of noradrenergic afferents and neurons

To determine the integrity of noradrenergic (NAergic) axons in forebrain, every 12^th^ brain sections were immunostained for Tyrosine Hydroxylase (TH). The length of TH+ afferents were estimated using the stereological length estimation with the spherical probes (Stereo Investigator; Micro Bright Field, Williston, VT, USA) as described previously.^14, 35^ Because of the regional variations in the densities of TH+ afferents, we focused our analysis on the selected subregions or ROIs (S1BF and dorsal hippocampus) for TH+ afferents.

To determine the integrity of the TH+ neurons in locus coeruleus (LC), every 4^th^ coronal sections through the hindbrain was immunostained for TH. The total number of TH+ neurons in the LC was determined from by counting neurons through the entire LC using the optical fractionator as described previously.^14, 35^ For unbiased stereological analysis of neuronal size (area and volume), we used the nucleator probe of the Stereo Investigator as described previously.^14, 35^

### Statistical analysis

One-way or two-way analysis of variance (ANOVA) was used to establish statistical significance using GraphPad Prism version 10. Tukey, Holm-Sidak, and Dunnett multiple comparison tests were applied as needed, and values of p < 0.05 were deemed statistically significant.

## ASSOCIATED CONTENT

### Supporting Information

The Supporting Information is available free of charge at https://pubs.acs.org/doi/XXXXXX.

- Brain levels of ψ-GSH after chronic treatment (Figure S1), effect of treatment on neuroinflammation in the hippocampus (Figure S2), effect of the prodrug on reactive astrocytes and microglia in the wild type mice (Figure S3), cytokine array analysis (Figure S4), effect of pro-ψ-GSH on atrophy of non-TH+ neurons (Figure S5), effect of pro-ψ-GSH on cortical TH+ afferents and atrophy (Figure S6), metabolite profile in prodrug treated mice (Figure S7), statistical analysis of cytokine expression (Table S1), biomarker analysis of hippocampal metabolites (table S2)

## AUTHOR INFORMATION

### Author Contributions

Conceptualization, S.S.M. and M.K.L.; experiments, S.P.R., A.I.F., W.X., S.P., K.C., and Y.Z.; data collection and analysis, S.P.R., A.I.F., W.X., S.P., K. C., M.K.L., and S.S.M.; writing— original draft preparation, S.P.R. and A.I.F..; writing—review and editing, S.S.M., M.K.L., R.V., E.L., W.X., and Y.Z.; supervision, S.S.M., R.V., and M.K.L.; funding acquisition, S.S.M. and M.K.L. The manuscript was written through contributions of all authors. All authors have given approval to the final version of the manuscript. ‡S.P.R. and A.I.F. contributed equally to this work.

## Funding

This research was funded by the National Institutes of Health grant to S.S.M. and M.K.L. (R01-AG062469), to M.K.L. (RF1-AG062135, R01-AG077743), and the Center for Drug Design (CDD), University of Minnesota.

## Competing interests

S.S.M. and R.V. are co-inventors on the patent applications relating to ψ-GSH and its analogs as treatment options for neurodegenerative disorders and liver diseases. The funders had no role in the design of the study; in the collection, analyses, or interpretation of data; in the writing of the manuscript; or in the decision to publish the results.

## Availability of data and materials

All data generated or analyzed during this study are included in this published article and in Supplementary data are available from the corresponding author on reasonable request.

## ACKNOWLEDGMENT

The authors thank Dr. Jiashu Xie for LC-MS/MS analysis of brain homogenates for quantification of methylglyoxal levels. We thank the Center for Metabolomics and Proteomics at the University of Minnesota for providing services related to metabolomics characterization of brain homogenates using Biocrates kit. We thank the Mouse Behavior Core at the University of Minnesota for equipment access and technical assistance. We also thank Joyce Meints for help with neuropathological analysis. The graphical abstract, the glyoxalase pathway, and timeline for animal experiments were created with BioRender.com.

## ABBREVIATIONS

Aβ: β-amyloid peptide
AD: Alzheimer’s disease
AGE: advanced glycation end products
GGT: γ-glutamyl transpeptidase
Glo-1: glyoxalase-1
GSH: glutathione
*i.c.v.*: intracerebroventricular
LC: locus coeruleus
MG: methylglyoxal
NAergic: noradrenergic
NOR: novel object recognition test
NTG: non-transgenic
PCA: principle component analysis
RAGE: receptor for advanced glycation end products
TBARS: thiobarbituric acid reactive substances.

