## Supporting Information for "Oral prodrug of a novel glutathione surrogate reverses metabolic dysregulation and attenuates neurodegenerative process in APP/PS1 mice"

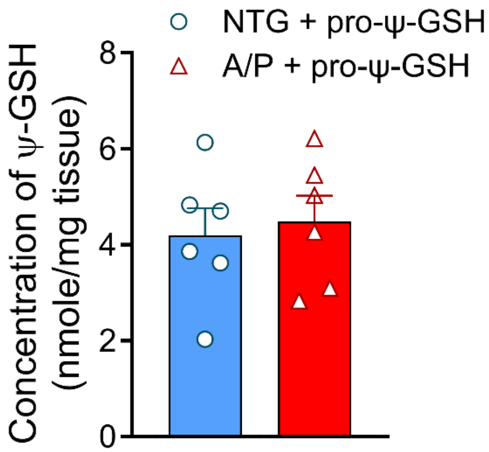


**Figure S1.** Measurement of brain levels of ψ-GSH after chronic administration of pro-ψ-GSH. Age-matched non-transgenic and APP/PS1 mice were treated with pro-ψ-GSH (250 mg/kg) orally for 12 weeks and total concentration in the brain was measured by LC-MS/MS. Brain levels of ψ-GSH were unaffected by genotype. Data are represented as the mean ± SEM.

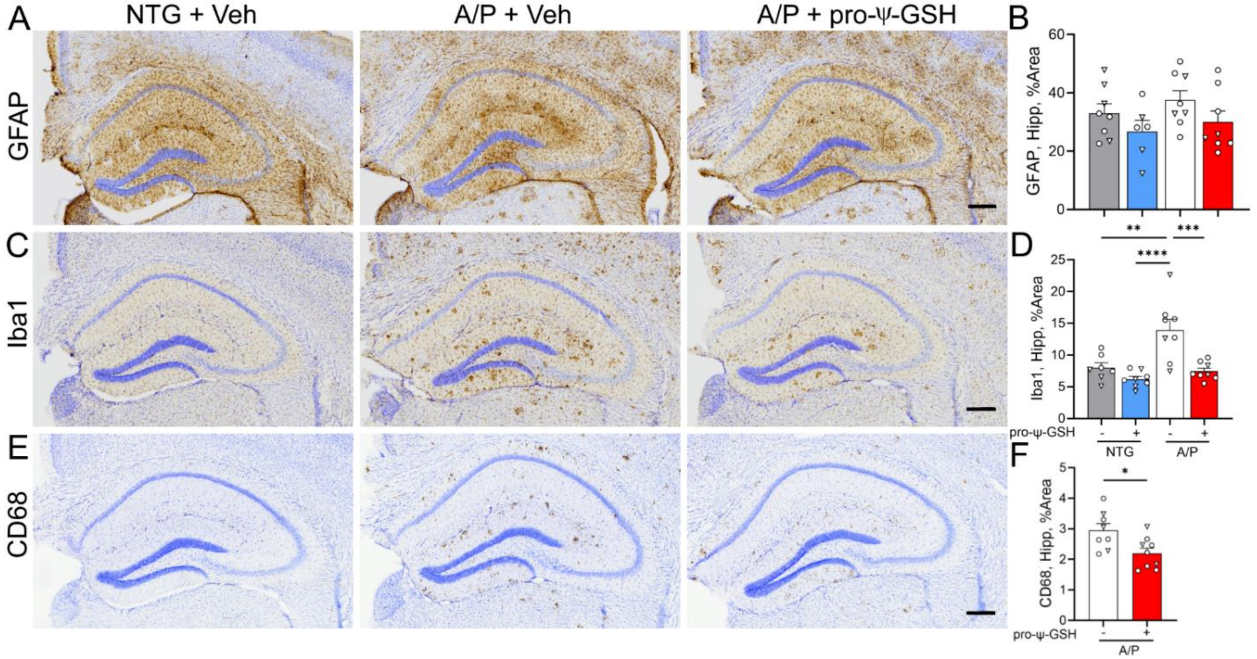


**Figure S2. Effect of oral pro-ψ-GSH treatment on hippocampal reactive astrocytosis and microglial activation in the APP/PS1 mouse model of AD.** (A, C, E) Representative images of brain sections stained for astrocytes (GFAP), total microglia (Iba1) and activated microglia (CD68) in hippocampal region of NTG and APPswe/PS1ΔE9 (A/P) mouse cohorts. Animals were treated with either Vehicle (Veh) or pro-ψ-GSH for 3 months prior to tissue collection. Scale bar, 250 µm. (B, D. F) Hippocampal glial reaction was quantified by defining the percent area covered by GFAP (B), Iba-1(D), or CD68 (F) immunoreactivity. The results show the expected increase in microglial activation in the Veh treated A/P mice compared to NTG mice. Consistent with reduced amyloid pathology, oral pro-ψ-GSH treatment significantly reduces microglial activation (Iba1 and CD68). Scale bar, 250 µm. Circles are males and triangles are females. Data are mean ± S.E.M.; One-way ANOVA with Tukey's post-hoc test. * $p < 0.05$, ** $p < 0.01$, *** $p < 0.005$, **** $p < 0.001$.

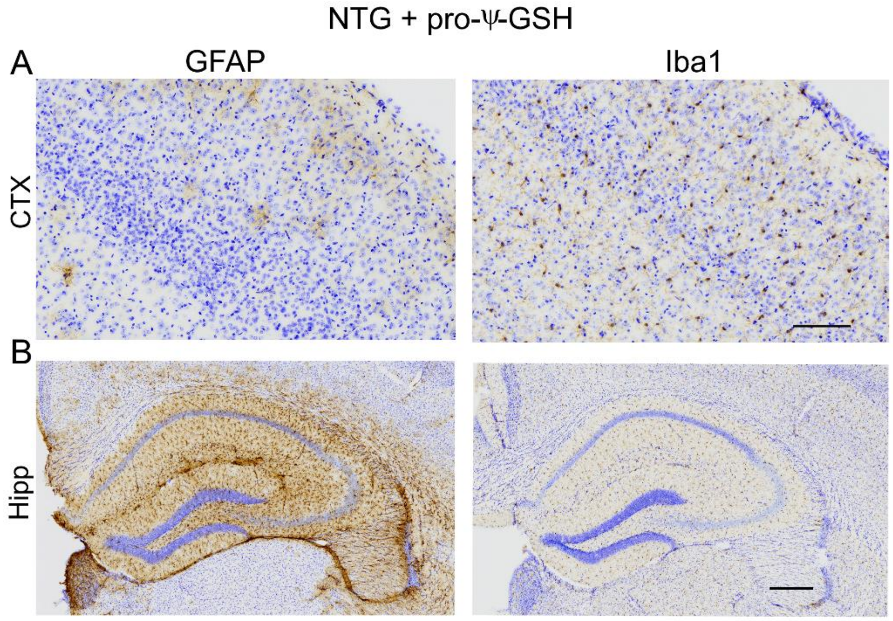


**Figure S3. Effect of oral pro-ψ-GSH treatment on reactive astrocytosis and microglial activation in cortex and hippocampal regions of NTG mice.** (A, B) Representative images of brain sections stained for astrocytes (GFAP) and total microglia (Iba1) in cortex and hippocampal region of NTG mice treated with pro-ψ-GSH. Scale bar, 100 µm for panel A; and 250 [illegible]M for panel B. The results show no increase in GFAP and Iba1 immunoreactivity compared to vehicle treated NTG controls, as shown in Figure 5.

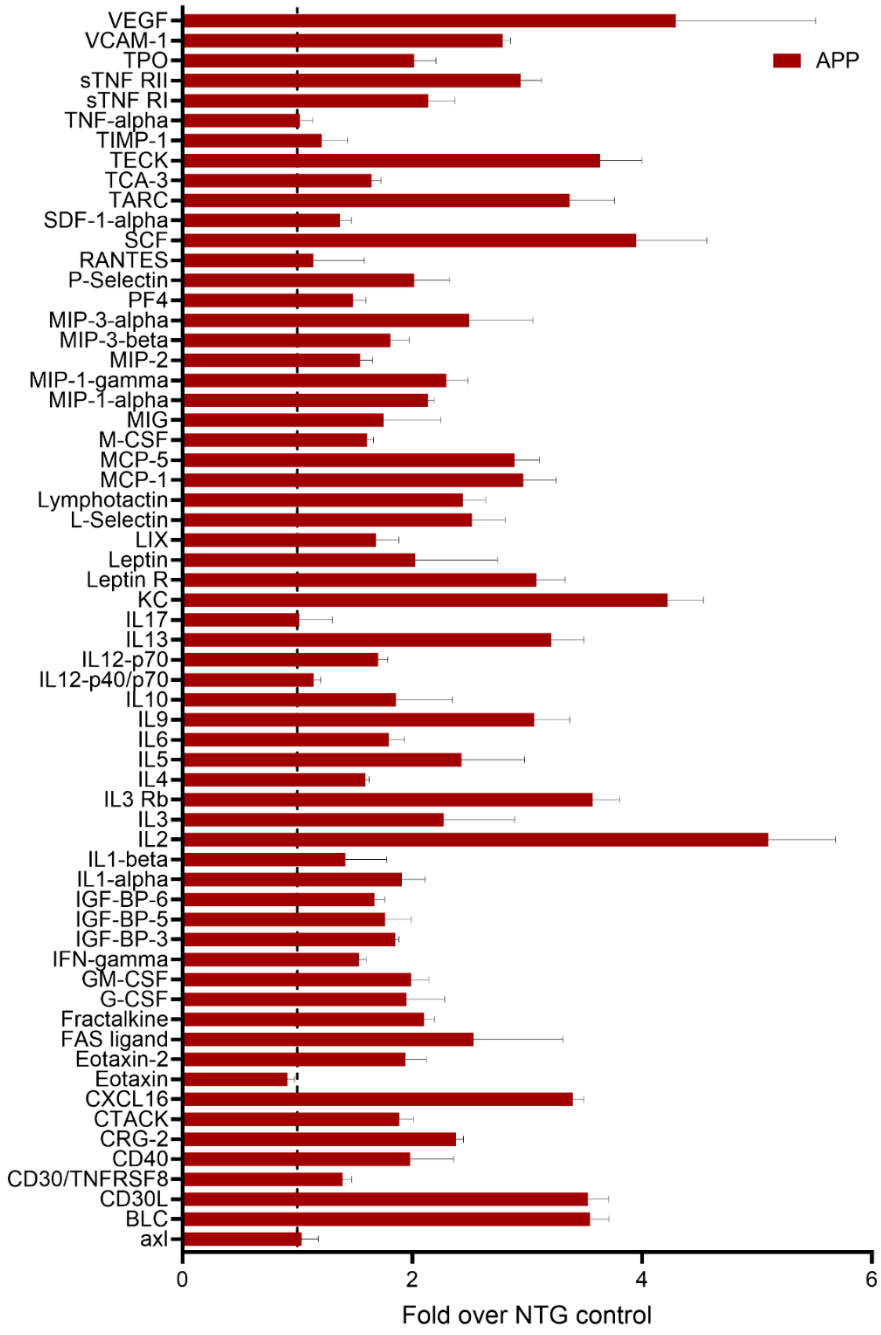


**Figure S4. Cytokine array analysis of mice based on genotype.** Changes in the inflammatory cytokine levels in hippocampal brain homogenates from APP/PS1 mice and age-matched non-transgenic littermates were analyzed as described in Methods. The data shown here is normalized with respect to NTG vehicle controls (dotted line at 1). Cytokine analysis showed a significant elevation of pro-inflammatory cytokines in saline treated APP/PS1 mice (N = 4 per group). Data are expressed as mean ± SEM.

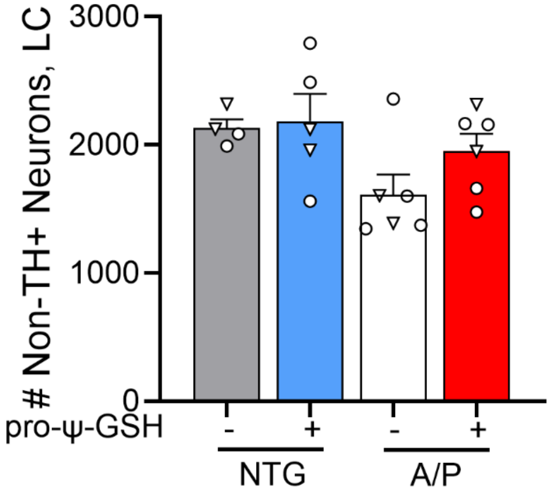


**Figure S5. Effect of oral pro-ψ-GSH treatment on atrophy of non-TH+ neurons in Locus Coeruleus (LC) in the APP/PS1 model.** Stereological counting of non-TH+ neurons in the LC do not show significant neuronal loss in A/P mice. Circles are males and triangles are females. Data are mean ± S.E.M. One-way ANOVA with Tukey's post-hoc test.

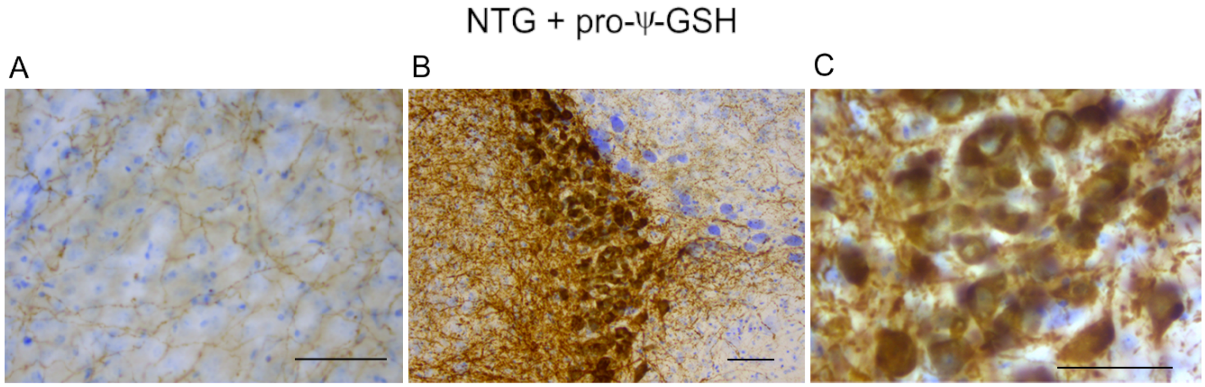


**Figure S6. Effect of oral pro-ψ-GSH treatment on cortical TH+ afferents and atrophy in Locus Coeruleus (LC) in NTG mice.** (A, B, C) Representative images of TH+ afferents (A) in S1BF and low (B) and high (C) magnification images of TH+ neurons of the LC of pro-ψ-GSH-treated NTG cohort. Quantitative analysis of TH+ afferent density show no effect of treatment when compared to NTG-vehicle subjects. Stereological counting of TH+ neurons in the LC do not show significant observation after treatment, as shown in Figure 7. Scale bar, 50 [illegible]m.

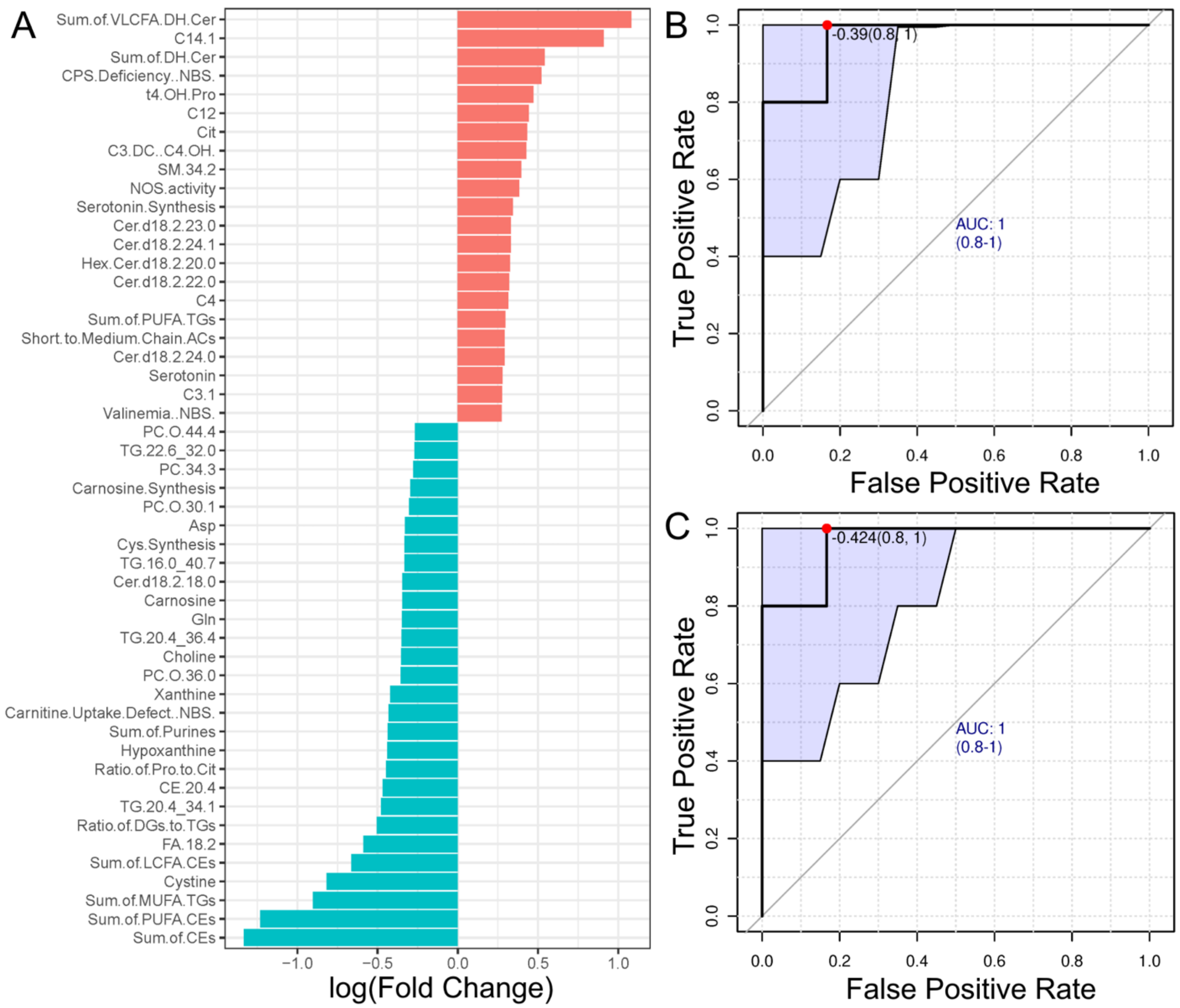


**Figure S7. Metabolite alterations caused by pro-ψ-GSH treatment in symptomatic APP/PS1 mice.** (A) Using pattern search function in Metaboanalyst, fold changes in metabolite concentrations between vehicle and prodrug treated APP/PS1 brains was plotted. Overall, the prodrug treatment reduced the levels of metabolites contributing toward disease pathology, while elevated the beneficial metabolites for neuronal function. (B-C) Biomarker analysis of hippocampal metabolites using ROC curve. PUFA-CEs (B) and sum of cholesteryl esters (CEs, C) were significantly modulated by pro-ψ-GSH and offered an AUC close to 1.

**Table S1. Two-way ANOVA analysis of cytokine expression difference between NTG + Vehicle and A/P + vehicle groups.**

| UNCORRECTED FISHER'S LSD | PREDICTED (LS) MEAN DIFF. | 95.00% CI OF DIFF. | PREDICTED (LS) MEAN DIFF. | BELOW THRESHOLD? | SUMMARY | INDIVIDUAL P VALUE |
|---|---|---|---|---|---|---|
| **AXL** | -0.0365 | -1.060 to 0.9873 | -0.0365 | No | ns | 0.944076 |
| **BLC** | -2.547 | -3.571 to -1.523 | -2.547 | Yes | * | 0.000002 |
| **CD30L** | -2.527 | -3.551 to -1.503 | -2.527 | Yes | * | 0.000002 |
| **CD30/TNFRSF8** | -0.394 | -1.418 to 0.6298 | -0.394 | No | ns | 0.449184 |
| **CD40** | -0.981 | -2.005 to 0.04279 | -0.981 | No | ns | 0.060295 |
| **CRG-2** | -1.38 | -2.403 to -0.3557 | -1.38 | Yes | * | 0.008472 |
| **CTACK** | -0.8875 | -1.911 to 0.1363 | -0.8875 | No | ns | 0.089004 |
| **CXCL16** | -2.397 | -3.421 to -1.373 | -2.397 | Yes | * | 0.000006 |
| **EOTAXIN** | 0.086 | -0.9378 to 1.110 | 0.086 | No | ns | 0.868727 |
| **EOTAXIN-2** | -0.9397 | -1.964 to 0.08404 | -0.9397 | No | ns | 0.071835 |
| **FAS LIGAND** | -1.533 | -2.557 to -0.5090 | -1.533 | Yes | * | 0.003496 |
| **FRACTALKINE** | -1.101 | -2.125 to -0.07746 | -1.101 | Yes | * | 0.03512 |
| **G-CSF** | -0.9485 | -1.972 to 0.07529 | -0.9485 | No | ns | 0.069246 |
| **GM-CSF** | -0.9892 | -2.013 to 0.03454 | -0.9892 | No | ns | 0.058183 |
| **IFN-GAMMA** | -0.5375 | -1.561 to 0.4863 | -0.5375 | No | ns | 0.302123 |
| **IGF-BP-3** | -0.852 | -1.876 to 0.1718 | -0.852 | No | ns | 0.102465 |
| **IGF-BP-5** | -0.7638 | -1.788 to 0.2600 | -0.7638 | No | ns | 0.143018 |
| **IGF-BP-6** | -0.671 | -1.695 to 0.3528 | -0.671 | No | ns | 0.197951 |
| **IL1-ALPHA** | -0.909 | -1.933 to 0.1148 | -0.909 | No | ns | 0.081574 |
| **IL1-BETA** | -0.4172 | -1.441 to 0.6065 | -0.4172 | No | ns | 0.422913 |
| **IL2** | -4.096 | -5.120 to -3.072 | -4.096 | Yes | * | <0.000001 |
| **IL3** | -1.273 | -2.297 to -0.2495 | -1.273 | Yes | * | 0.014999 |
| **IL3 RB** | -2.57 | -3.594 to -1.546 | -2.57 | Yes | * | 0.000001 |
| **IL4** | -0.5935 | -1.617 to 0.4303 | -0.5935 | No | ns | 0.254649 |
| **IL5** | -1.428 | -2.451 to -0.4037 | -1.428 | Yes | * | 0.00647 |
| **IL6** | -0.795 | -1.819 to 0.2288 | -0.795 | No | ns | 0.127434 |
| **IL9** | -2.061 | -3.085 to -1.037 | -2.061 | Yes | * | 0.000096 |
| **IL10** | -0.8565 | -1.880 to 0.1673 | -0.8565 | No | ns | 0.100673 |
| **IL12-P40/P70** | -0.142 | -1.166 to 0.8818 | -0.142 | No | ns | 0.784942 |
| **IL12-P70** | -0.703 | -1.727 to 0.3208 | -0.703 | No | ns | 0.177469 |
| **IL13** | -2.208 | -3.232 to -1.184 | -2.208 | Yes | * | 0.000031 |

| | | | | | | |
|---|---|---|---|---|---|---|
| **IL17** | -0.0175 | -1.041 to 1.006 | -0.0175 | No | ns | 0.97317 |
| **KC** | -3.221 | -4.244 to -2.197 | -3.221 | Yes | * | <0.000001 |
| **LEPTIN R** | -2.08 | -3.104 to -1.056 | -2.08 | Yes | * | 0.000083 |
| **LEPTIN** | -1.026 | -2.050 to -0.002206 | -1.026 | Yes | * | 0.049512 |
| **LIX** | -0.683 | -1.707 to 0.3408 | -0.683 | No | ns | 0.190075 |
| **L-SELECTIN** | -1.52 | -2.544 to -0.4960 | -1.52 | Yes | * | 0.003779 |
| **LYMPHOTACTIN** | -1.44 | -2.464 to -0.4162 | -1.44 | Yes | * | 0.006025 |
| **MCP-1** | -1.967 | -2.991 to -0.9430 | -1.967 | Yes | * | 0.000194 |
| **MCP-5** | -1.89 | -2.914 to -0.8660 | -1.89 | Yes | * | 0.000337 |
| **M-CSF** | -0.6065 | -1.630 to 0.4173 | -0.6065 | No | ns | 0.244418 |
| **MIG** | -0.749 | -1.773 to 0.2748 | -0.749 | No | ns | 0.150867 |
| **MIP-1-ALPHA** | -1.137 | -2.161 to -0.1130 | -1.137 | Yes | * | 0.029686 |
| **MIP-1-GAMMA** | -1.295 | -2.319 to -0.2712 | -1.295 | Yes | * | 0.013381 |
| **MIP-2** | -0.5465 | -1.570 to 0.4773 | -0.5465 | No | ns | 0.294118 |
| **MIP-3-BETA** | -0.8112 | -1.835 to 0.2125 | -0.8112 | No | ns | 0.119874 |
| **MIP-3-ALPHA** | -1.496 | -2.519 to -0.4717 | -1.496 | Yes | * | 0.004364 |
| **PF4** | -0.4838 | -1.508 to 0.5400 | -0.4838 | No | ns | 0.352946 |
| **P-SELECTIN** | -1.015 | -2.039 to 0.008544 | -1.015 | No | ns | 0.051928 |
| **RANTES** | -0.1385 | -1.162 to 0.8853 | -0.1385 | No | ns | 0.790117 |
| **SCF** | -2.949 | -3.973 to -1.925 | -2.949 | Yes | * | <0.000001 |
| **SDF-1-ALPHA** | -0.37 | -1.394 to 0.6538 | -0.37 | No | ns | 0.477254 |
| **TARC** | -2.369 | -3.393 to -1.345 | -2.369 | Yes | * | 0.000008 |
| **TCA-3** | -0.6453 | -1.669 to 0.3785 | -0.6453 | No | ns | 0.215656 |
| **TECK** | -2.635 | -3.659 to -1.611 | -2.635 | Yes | * | <0.000001 |
| **TIMP-1** | -0.213 | -1.237 to 0.8108 | -0.213 | No | ns | 0.682329 |
| **TNF-ALPHA** | -0.0215 | -1.045 to 1.002 | -0.0215 | No | ns | 0.967041 |
| **STNF RI** | -1.139 | -2.163 to -0.1152 | -1.139 | Yes | * | 0.029367 |
| **STNF RII** | -1.942 | -2.966 to -0.9185 | -1.942 | Yes | * | 0.000232 |
| **TPO** | -1.015 | -2.039 to 0.008544 | -1.015 | No | ns | 0.051928 |
| **VCAM-1** | -1.787 | -2.811 to -0.7632 | -1.787 | Yes | * | 0.000688 |
| **VEGF** | -3.293 | -4.317 to -2.269 | -3.293 | Yes | * | <0.000001 |

**Table S2. Biomarker analysis of hippocampal metabolites using ROC curve to determine pro-ψ-GSH treatment-specific effects difference in the APP/PS1 model.**

| | AUC | P value | FC | clusters |
|---|---|---|---|---|
| **PC.34.2** | 0.9667 | 0.0015 | -0.2156 | 3 |
| **TG.20.4_34.1** | 0.9667 | 0.0043 | -0.4772 | 3 |
| **PC.34.3** | 0.9333 | 0.0062 | -0.2779 | 3 |
| **Sum.of.PUFA.CEs** | 0.9667 | 0.0084 | -1.2320 | 3 |
| **Sum.of.CEs** | 0.9667 | 0.0085 | -1.3330 | 3 |
| **C12** | 0.9333 | 0.0096 | 0.4430 | 1 |
| **C14.1** | 0.9333 | 0.0105 | 0.9079 | 1 |
| **PC.O.30.0** | 0.8500 | 0.0268 | -0.1787 | 2 |
| **C3.1** | 0.8667 | 0.0278 | 0.2748 | 1 |
| **PC.42.6** | 0.8667 | 0.0324 | -0.1641 | 3 |
| **Ala** | 0.8500 | 0.0332 | 0.2208 | 1 |
| **C16.1** | 0.8667 | 0.0345 | 0.2644 | 1 |
| **PC.O.44.6** | 0.8667 | 0.0379 | -0.2649 | 3 |
| **PC.O.30.1** | 0.9167 | 0.0392 | -0.3023 | 2 |
| **Cer.d18.2.24.1** | 0.8333 | 0.0419 | 0.329 | 5 |
| **Creatinine** | 0.7667 | 0.0458 | 0.1820 | 1 |
| **TG.20.4_36.4** | 0.8667 | 0.0462 | -0.3502 | 3 |
| **PC.36.6** | 0.9333 | 0.0488 | -0.1775 | 3 |
| **SM.36.2** | 0.8500 | 0.0504 | 0.2579 | 5 |
